# Synergizing algorithmic design, photoclick chemistry and multi-material volumetric printing for accelerating complex shape engineering

**DOI:** 10.1101/2022.11.29.518318

**Authors:** Parth Chansoria, Dominic Rütsche, Anny Wang, Hao Liu, Davide D’Angella, Riccardo Rizzo, Amelia Hasenauer, Patrick Weber, Nafeesah Bte Mohamed Ibrahim, Nina Korshunova, Marcy Zenobi-Wong

## Abstract

Accelerating the designing and manufacturing of complex shapes has been a driving factor of modern industrialization. This has led to numerous advances in computational design and modeling and novel additive manufacturing (AM) techniques that can create complex shapes for bespoke applications. By combining a new coding-based design approach with high-throughput volumetric printing, we envision a new approach to transform the way we design and fabricate complex shapes. Here, we demonstrate an algorithmic voxel-based approach, which can rapidly generate and analyze porous structures, auxetic meshes and cylinders, or perfusable constructs. We use this design scheme in conjunction with new approaches for multi-material volumetric printing based on thiol-ene photoclick chemistry to rapidly fabricate complex heterogeneous structures. Collectively, the new design and fabrication technique we demonstrate can be used across a wide-spectrum of products such as actuators, biomedical implants and grafts, or tissue and disease models.

**Teaser:** A new scheme of rapidly designing and printing complex multi-material structures for implant and tissue graft applications.

## Introduction

Designing and manufacturing complex shapes at increased throughput has been a key challenge of modern engineering. This is highly relevant in the field of biomedical implants and grafts. Using advanced computer-aided design (CAD) and computational modeling (CM) tools, complex implants are being developed which offer better mechanics (reduced weight and increased load-bearing capability, etc.) or patient safety and comfort, as replacements of simpler implants. For example, the designs of arterial stents have transitioned from the conventional porous shapes to auxetic architectures (*1, 2*), which allow easy radial expansion of the stents while reducing the risk of stent malapposition and foreshortening (*1*). Such auxetic shapes have also recently paved the way to a new range of patches and tissue grafts for regenerative applications such as those repairing cardiac (*3*) and pulmonary pathologies (*4*), where auxetic patches allow easy conformation to organ deformation and outperform non-auxetic patches (*5*). In addition, auxetic structures are increasingly being used as actuators (*6, 7*) or load-bearing structures (*8, 9*). Furthermore, advanced CAD and CM tools are increasingly being used in tissue engineering to design complex perfusable structures, such as perfusable vascularized biomimetic tissue models for studying biogenesis and disease progression and treatment (*10, 11*). Unfortunately, to this date, CAD and CM still largely require manually defining the complex geometrical relationships and boundary conditions. Furthermore, one needs to analyze a wide array of design iterations to derive the optimized design for the application. For example, auxetic patches developed for different dynamic organs (lung, heart, etc.) need to conform to the stiffness and Poisson’s ratios of the different organs, and may require analyzing hundreds of design iterations and their computational modeling to find the right patch design for an organ (*4*). Herein, we present a new algorithmic approach to designing complex shapes within seconds, which can rapidly generate a large array of design iterations within minutes.

To fabricate the complex shapes, additive manufacturing techniques involving layer-by-layer material deposition offer a wide range of achievable resolution (typically 10 – 500 µm) and throughput (0.01 – 1000 mm^3^/hr). Recently, volumetric printing (VP), also termed as volumetric additive manufacturing, has emerged as a powerful technique towards the fabrication of high-resolution structures (up to 100 µm) within tens of seconds. VP relies on computed axial lithography, where the vial containing photocrosslinkable resin (photoresin) is rotated with dynamically evolving light patterns (images) projected into the resin (*12–14*). The superposition of the projected images leads to a spatially localized increase in the free radicals produced from the photoinitiator, which induces crosslinking of the photoresin into the desired shape. The photo-rheology of the resins used in VP typically depicts a non-linear response, where the crosslinking is induced after a threshold of light dose is achieved. There are also non-rotational methods involving image projections from multiple sides (typically front, side and bottom) to generate a spatially localized increase in light dose within the photocrosslinkable matrix, which induces crosslinking of the material into the desired shape (*15, 16*). Compared to multi-direction projections, computed tomography leads to better resolution and shape fidelity of the printed construct as the image can be changed continuously (*12*). We recently demonstrated that the tomographic printing duration can be further reduced to only a few seconds by using thiol-ene photoclick chemistry-based resins (*17*). Herein, insensitivity to oxygen and a homogeneous network formation within the step-growth polymerized matrices can reduce internal structural stresses and shrinkage after printing compared to constructs resulting from chain-growth polymerization (*17, 18*). Further, refractive index (RI) matching (*19*) and fine-tuning of light dose (*20*) has enabled high resolution printing while allowing encapsulation of high density of cells and organoids (*19*), which are critical to biomimetic tissue engineering. In an attempt to set a roadmap for high throughput design and fabrication of biomedical implants and grafts, we demonstrate how the complex algorithmically designed structures can be rapidly synthesized using photoclickable gelatin-based matrices within VP.

Furthermore, applications of VP have been limited to constructs based on single material compositions, and there is a critical need for new methods to fabricate multi-material structures, which can widen the applicability of VP into biomimetic structures as well as tissue and disease models. In this work, we demonstrate new approaches for multimaterial VP (Multimat VP). We print selected architectures with different photoresin compositions along the length or the thickness of the constructs. Further we demonstrate how the designs can be wrapped around more complex objects such as a heart, which paves the way for rapidly printing organ-specific auxetic meshes. Finally, we also highlight how complex multimaterial perfusable architectures such as alveoli can be rapidly designed and fabricated using our approach. The synergy of algorithmic design, photoclick chemistry and Multimat VP (**Figure 1**) offers a transformational approach to rapidly designing and fabricating complex multimaterial shapes.

**Figure 1.**
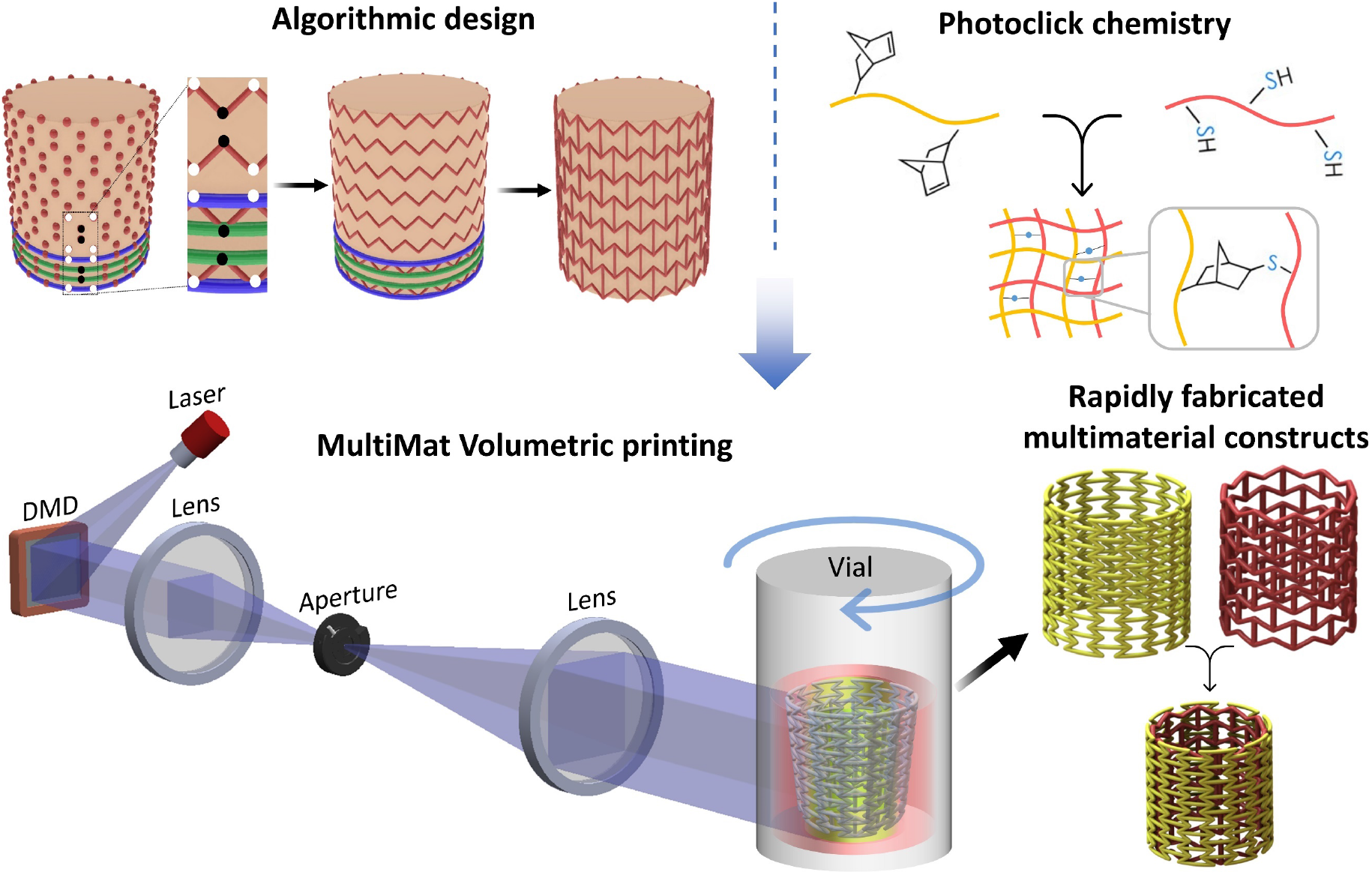
Schematic of the proposed concept to rapidly design and fabricate complex structures. Algorithmic design is used to rapidly create large arrays of design iterations, photoclick chemistry-based bioresins allow rapid fabrication of constructs, and multi-material volumetric printing (Multimat VP) approach rapidly fabricates complex constructs made from heterogenous resin compositions.

## Results

### Rapid algorithmic design and multimaterial volumetric printing of auxetic meshes

Auxetic shapes serve as an ideal template for algorithmic design as the structural interrelationships can be defined via governing equations. 2D auxetic meshes have been increasingly utilized as patches which feature tailorable negative Poisson’s ratios and directional stiffness to easily conform to dynamic organs such as the lung (*4*) or the heart (*3, 21*). **Figure 2A** illustrates the algorithmic design scheme for auxetic meshes featuring re-entrant honeycomb, sinusoidal ligaments, arrowhead and pinwheel meshes. All of these meshes feature different stiffness and Poisson’s ratios, which can be selectively matched to different dynamic organs (e.g., lung, heart, stomach, bladder, etc.) (*4, 5*). The equations governing the design algorithms have been provided in the Supplemental Information. Briefly, for re-entrant honeycomb meshes, vertical baselines containing the vertices of the mesh elements are established at a defined separation, followed by offsetting the position of every alternate vertex either ahead (positive offset) or behind (negative offset) the baseline. Herein, every consecutive baseline has opposing offsetting of the vertices (i.e., if the vertex at any baseline has a positive offset from the vertex, then the consecutive baseline will have a negative offset from the vertex). Next, the vertices are connected along the vertical direction, followed by connecting alternate vertices which are at positive and negative offset, along the horizontal direction. For the arrowhead architectures, the meshes are offset such that each baseline features the same offset pattern of the vertices. The vertices are then connected such that the negative offset vertices of any baseline are connected to the positive offset vertices along the consecutive baseline. For creating the sinusoidal ligament meshes, sine functions are created with period length (2π) spanning two baselines, and the sinusoidal mesh in every other baseline is shifted by a phase spanning the distance between two consecutive baselines. The pinwheel meshes are fabricated in a similar way, except there is no offset between the sinusoids across the baselines. In **Figure 2B**, we present select architectures created by changing dimensions of the auxetic design features. Using this algorithmic approach, several hundred iterations of any auxetic mesh can be rapidly created (average compilation time per design on a personal computer at 1.3 GHz and 32 GB RAM was 0.04 s).

**Figure 2.**
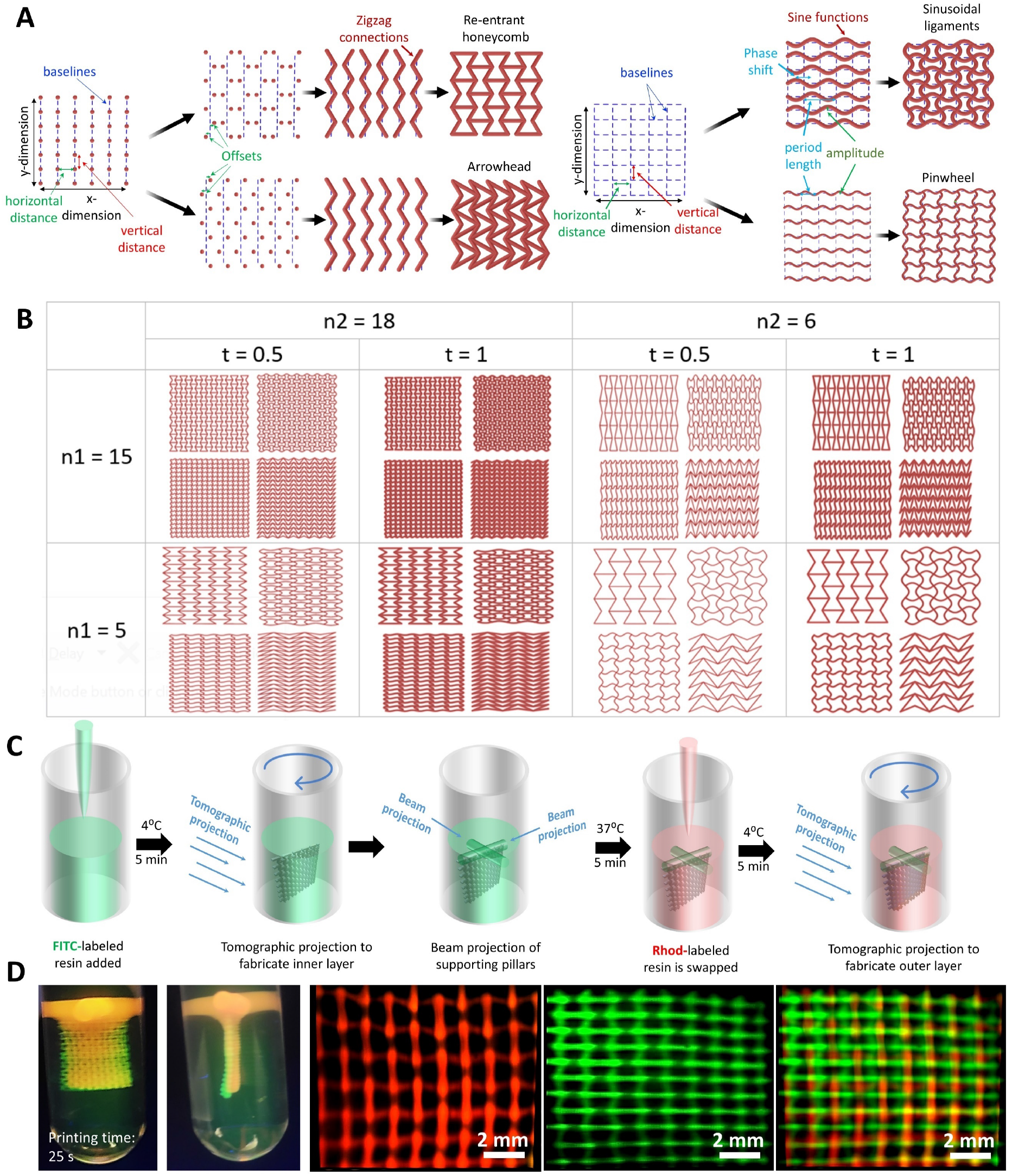
Rapid design and fabrication of auxetic meshes. **A.** Design rationale for creating different auxetic meshes – Re-entrant honeycomb, arrowhead, sinusoidal ligaments and pinwheel (see **Supplemental Information** for governing equations). **B**. Rapidly generated design iterations for the auxetic architectures by varying select design parameters (n1, n2 and t represent the number of vertical and horizontal elements, and thickness of the elements, respectively). **C**. Scheme of fabrication of the multimaterial auxetic meshes featuring different designs and resin compositions along the thickness. **D**. Rapidly fabricated (25 s printing time) auxetic meshes featuring horizontally and vertically oriented re-entrant auxetic meshes made of Rhodamine (Rhod)-labeled, FITC-labeled GelNB/GelSH resins, respectively, along the thickness. Images of the meshes have been captured using light sheet microscopy.

With auxetic meshes as the template, we present the first scheme for the rapid Multimat VP (**Figure 2C**). Here, the inner mesh is created using tomographic projections in a vial filled with thermo-reversibly crosslinked rhodamine-labeled fluorescent resin containing norbornene-modified gelatin (GelNB) and thiolated gelatin (GelSH) at 5% w/v (total gelatin content) in phosphate buffered saline (PBS) (see dose optimization and resolution tests for different photoclick materials in **Figure S1**). After the first tomographic projection, the mesh is localized in the resin container by projecting supporting beams (<u>created by projecting a circular image (Φ = 3 mm) for 5 s without rotating the vial</u>) at the top of the mesh. This prevents the printed structure from falling when the vial is heated up to 37°C to remove the non-photocrosslinked resin. The resin in the container is then interchanged with a FITC (fluorescein isothiocyanate)-labeled resin, followed by tomographic projections of the outer mesh. Addition of either Rhodamine or FITC did not affect the absorption of light at 405 nm and the RI of the resin (RI = 1.34 for GelNB/GelSH), thereby not affecting the light path during tomographic projections. **Figure 2D** demonstrates the rapidly fabricated (25 s total printing time) bi-layered auxetic mesh comprising of different resin compositions (Rhod-labeled and FITC-labeled GelNB/GelSH) and vertically and horizontally oriented re-entrant honeycomb meshes in the first and second layers, respectively. The images have been captured using light sheet microscopy (*22*), and the supporting pillars facilitate image capturing by stabilizing the constructs within the printing vials. Notably, post photocrosslinking, the RI of the GelNB/GelSH resin increases by ∼0.002 (i.e., RI = 1.342), which, as per our observations, did not critically affect the feature dimensions. The minimum feature size of the second layer (∼ 275 µm) was within ±5% as that of the first layer (∼ 264 µm).

Of note, the supporting pillars in the first Multimat VP scheme need to be removed post printing. The second printing scheme for Multimat VP does not require projection of supporting pillars. This scheme is demonstrated in **Figure 3A**. The scheme utilizes filling the first resin in the printing vial, followed by adding the second resin. Here, the two resin compositions are prevented from mixing into each other through thermo-reversible crosslinking of each resin formulation at 4°C prior to adding the subsequent one. Alternatively, a high viscosity resin could also be used to prevent mixing of the resins during the short printing duration. After the different resins are added, the entire construct is printed at once via tomographic projections (printing time 12 s). We use this Multimat VP technique to fabricate a re-entrant honeycomb mesh with vertically oriented elements featuring FITC and Rhod-labeled GelNB/GelSH resin in the bottom and top portions, respectively (**Figure 3B**). Of note, the supporting beams are still added to the construct to suspend it within the printing vial, which facilitates its imaging using light sheet microscopy. Here, gelatin-based resin has been used since it allows the layers to thermo-reversibly crosslink when the temperature is reduced, thereby preventing the photocrosslinked structures from falling under their own weight due to gravity. The gelatin could also be replaced with other materials such as pluronic or hyaluronic acid, etc., provided they increase the viscosity substantially to prevent the structures from displacing during the short printing duration. This Multimat VP approach can be used to print tissue interfaces. As an example, we printed the same auxetic mesh with GelNB/GelSH across the top and bottom, but the top compartment consisted of myoblasts (C2C12 murine), and the bottom compartment consisted of fibroblasts (3T3 murine), labelled with cell tracker red and green, respectively. After 4 weeks of maturation, the top compartment selectively exhibited myo-heavy chain staining, while both compartments featured collagen I staining (**Figure 3D**).

**Figure 3.**
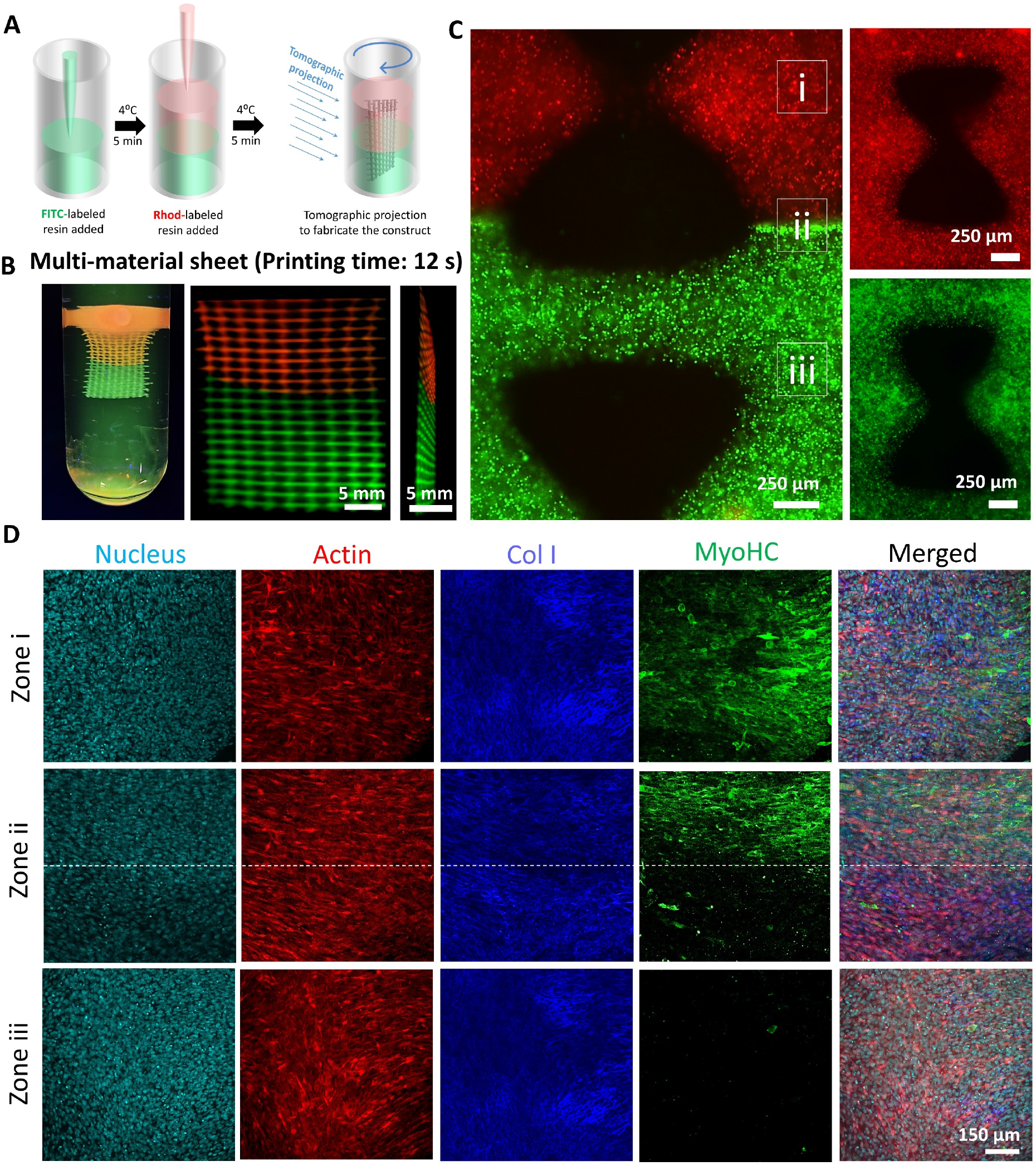
Fabrication of auxetic meshes featuring different resin compositions across the length of the meshes. **A**. Scheme of filling different resin compositions and printing the entire construct at once. **B**. Printed constructs featuring Rhod-labeled or FITC-labeled GelNB/GelSH matrix across the two layers. **C**. Micrographs of muscle-connective tissue model made of GelNB/GelSH resin. The top and bottom portions comprise of C2C12 myoblasts labeled with cell tracker red and 3T3 fibroblasts labeled with cell tracker green, respectively. **D**. After 4-week long culture in DMEM containing 2% w/v horse serum, the muscle-mimicking mesh containing C2C12 cells (Zone i) exhibits Myosin heavy chain (MyoHC) staining. The interface at Zone ii shows the distinction in MyoHC staining across the two layers, whereas the connective tissue-mimicking mesh containing 3T3 cells (Zone iii) does not demonstrate MyoHC staining.

### Rapid algorithmic design and multimaterial volumetric printing of auxetic cylinders

Cylindrical auxetic meshes, such as those increasingly being used for fabrication of stents (*1, 2*) can be even more complicated to design when compared to 2D meshes, especially when the design elements need to form a continuum across a cylindrical contour. Here, creating design iterations of the auxetic cylinders is particularly challenging. **Figure 4A** illustrates the algorithmic design scheme of cylindrical auxetic meshes featuring re-entrant honeycomb and sinusoidal ligament elements (detailed equations have been provided in the **Supplemental Information**). The design schemes of auxetic cylinders featuring arrowhead or pinwheel architectures have been provided in the **Supplemental Information** and **Figure S2**. For creating any auxetic mesh, we first convert the cartesian coordinate system to a cylindrical coordinate system to be able to create a cylinder and define points on it. For auxetic cylinders featuring vertically-oriented re-entrant honeycomb lattices, we create vertical zigzag lines by defining the coordinates of all corner points alternating on two sides of a base line (**Figure 4A**). The alternate corner points lie along parallel circles with a phase shift commensurate with the width of the re-entrant meshes. After all the corner points of the vertical zigzag lines are defined, these are connected with straight beams. To match the cylinder curvature, every point on the initial straight beam is translated onto the cylinder using a wrapping algorithm (details provided in the **Supplemental Information**). Finally, to create the horizontal beams, every second pair of neighboring points on each ring of the cylinder is connected by an arc using the same wrapping method, with the positions of the horizontal beams alternating along the axis of the cylinder. For auxetic cylinders featuring horizontally oriented re-entrant honeycomb meshes, the corner points of each zigzag ring are composed of two circles consisting of points with a certain phase shift and angular step to each other. After defining all corner points of the zigzag rings, these can be connected. Before generating the connecting beams, every beam between two corner points is translated onto the cylinder coat using the wrapping method used previously. Finally, vertical beams are constructed between neighboring zigzag rings. For the auxetic cylinders with sinusoidal ligament meshes, sinusoidal functions are created along the cylinder, with consecutive functions featuring a phase shift with the previous sinusoid. Next, a vertical sine wave is generated to create the auxetic cylinders with sinusoidal elements. The starting and ending points of a vertical line are on the crossing points of the bottom and top circles and their baselines respectively. Similar to the 2D auxetic meshes, this algorithmic design scheme allows us to create a wide array of cylinders within a matter of seconds (**Figure 4B** and **Figure S2**).

**Figure 4.**
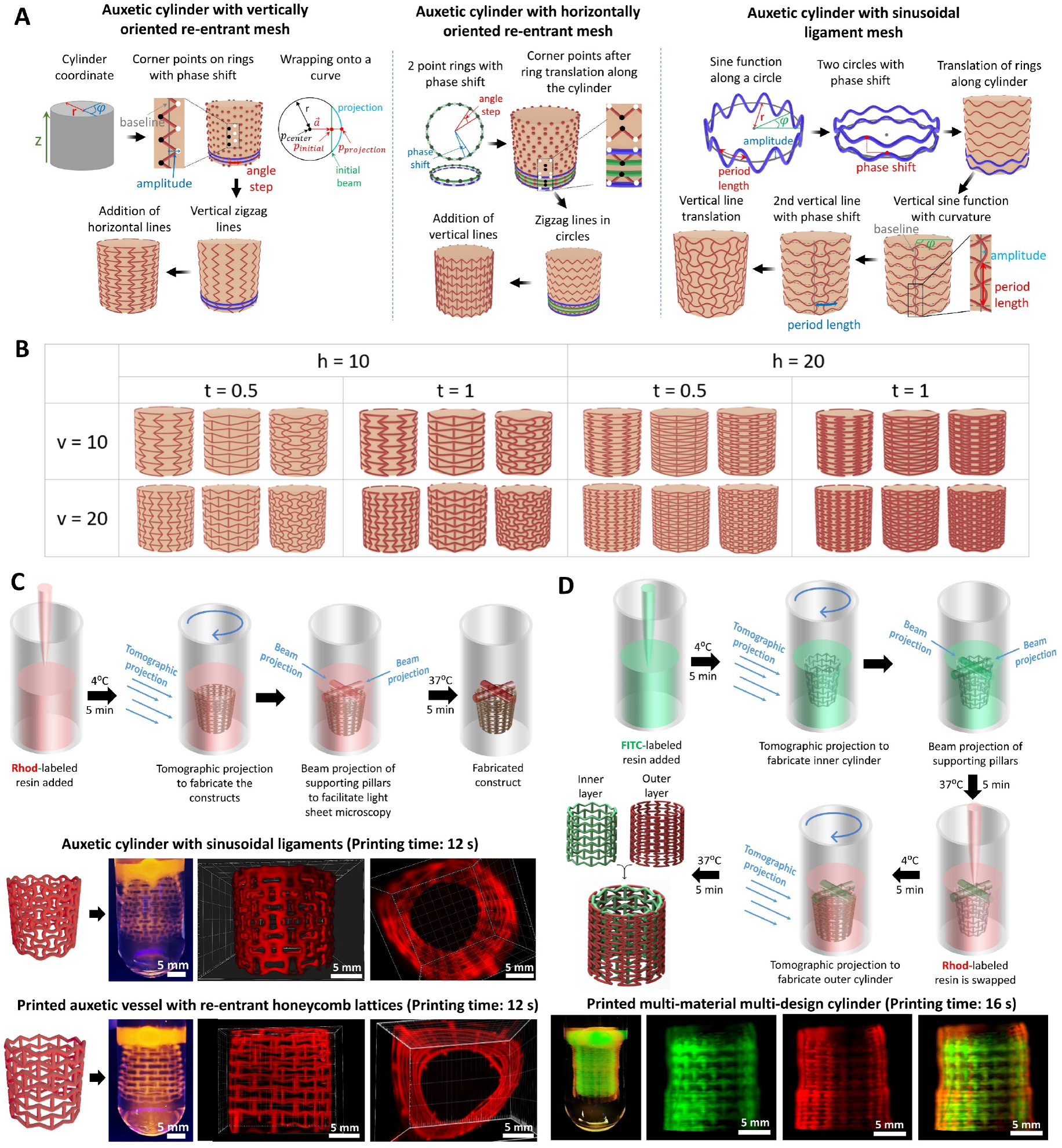
Rapid design and fabrication of auxetic cylindrical meshes. **A**. Algorithmic design rationales for the fabrication of auxetic cylindrical constructs featuring re-entrant and sinusoidal ligament meshes (see **Supplemental Information** for governing equations). **B**. Design arrays featuring selected iterations of the design parameters (h, t, and v represent the number of horizontal lines from top to bottom, thickness of the elements, number of vertical lines in the ring, respectively). **C**. Rapid fabrication of select architectures using volumetric printing, and compiled micrographs of the printed architectures (images are captured using the light sheet microscopy). **D**. Resin swapping scheme for the fabrication of bi-layered cylindrical auxetic meshes and printed meshes demonstrating horizontally and vertically oriented re-entrant honeycomb architectures across the inner and outer layers, respectively.

These auxetic architectures can also be printed within 12 s using photoclick resins within VP (**Figure 4C**). Using the first Multimat VP strategy of using support pillars to allow the construct to be suspended in the printing vial, which allows changing of the resin and re-printing, we also demonstrate how multimaterial and multidesign auxetic cylindrical meshes can be fabricated. **Figure 4D** demonstrates auxetic cylindrical meshes made of FITC-labeled GelNB/GelSH in the shape of horizontally-oriented re-entrant honeycomb mesh in the inner layer, and Rhod-labeled GelNB/GelSH in the shape of vertically-oriented re-entrant honeycomb mesh in the outer layer, respectively.

### Algorithmically-defined computational models for rapid screening of structural mechanics

Similar to algorithmic design, we show that algorithmic computational modeling can also substantially accelerate the process of determining the mechanics of a wide array of complex structures. Here we use the auxetic meshes and cylinders as the templates for our finite cell method (*23*)-based computational models. As opposed to the conventional approaches, the embedded simulation approach eliminates the necessity of the labor-intensive procedure for meshing and defining of boundary constraints. Instead, the geometry is embedded in a regular grid with vanishing stiffness. The physical model is then recovered by applying a voxel-based integration rule (*24*). Using this methodology allowed us to computationally model the auxetic meshes and cylinders at an average duration of approximately 4.5 s and 30 s, respectively, in a 1.3 GHz personal computer with 32 GB RAM. As a result, we were able to quickly derive the Poisson’s ratios of thousands of auxetic structures without any manual intervention. The corresponding results have been shown in **Figure 5**. The deformation of selected meshes have been shown in **Figure 5A**. Of these, the re-entrant meshes demonstrate the widest range of Poisson’s ratios (from 0.1 to - 10.9, **Figure 5B**) based on the different combinations of the design features (n1, n2 and t). In contrast to the auxetic meshes, the auxetic designs for cylinders (selected outputs are shown in **Figure 5C**) did not demonstrate negative Poisson’s ratios, but a wide range of positive Poisson’s ratios (**Figure 5D**). This means that the cylinders are actually contracting when they are stretched radially, even though the meshes feature auxetic designs. Here, the horizontally-oriented auxetic cylinders demonstrate Poisson’s ratios varying from 0.2 to 1.1 based on different combinations of the design features (h, v and t). Such a wide range of Poisson’s ratios for the different meshes and cylinders offers unique applications such as in actuators (*6, 7*), large structural components (*8, 9*), implants (*25, 26*) or organ-specific patches (*3, 4*).

**Figure 5.**
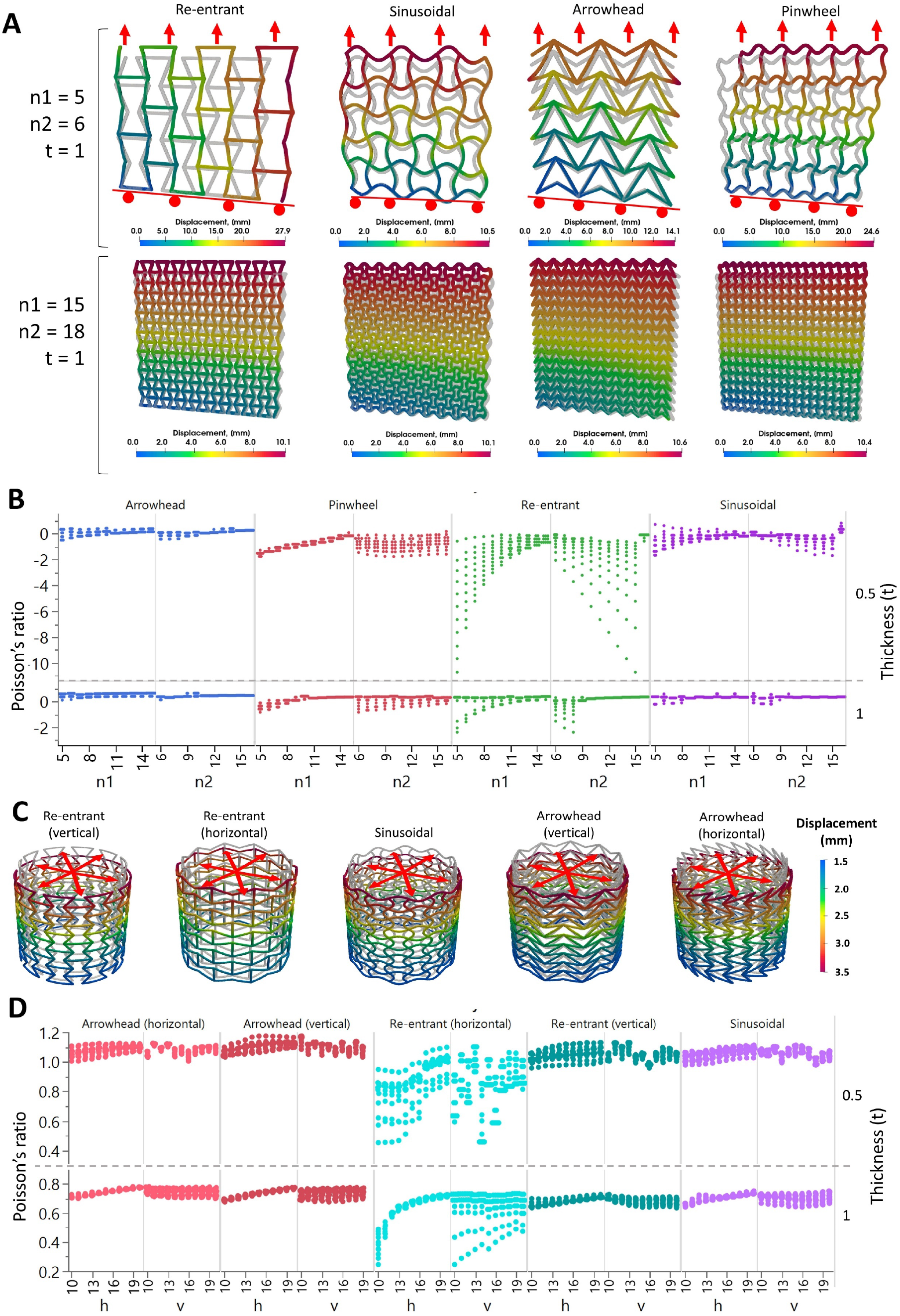
Algorithmically-derived computational estimates of structural mechanics (original mesh or cylinder is shown in grey). **A**. Selected computational outcomes (n1, n2 and t represent the number of vertical and horizontal elements, and thickness of the elements, respectively) demonstrating the negative Poisson’s ratios of the structures (i.e., structures expand laterally when stretched longitudinally (Note: A roller constraint was applied on the bottom of the mesh to allow free lateral expansion). **B**. Computational estimates of the Poisson’s ratios of the different auxetic meshes (244 designs were analyzed per auxetic mesh), based on different combinations of n1, n2 and t. Of note: The thickness has been plotted as a second y axis on the right. **C**. Selected computational outcomes of the auxetic cylinders. **D**. Computational estimates of the Poisson’s ratios of the cylinders based on the different design parameters (h, t, and v represent the number of horizontal lines from top to bottom, thickness of the elements, number of vertical lines in the ring, respectively; 1187 iterations were analyzed per cylinder). Thickness (t) is plotted as a second y axis on the right

### Organ-specific auxetic meshes

To demonstrate the level of design complexity that the algorithmic schemes can address, we demonstrate how the auxetic meshes can be wrapped around more complex shapes such as the heart (**Figure 6A)**. Here, we start with a square 2D auxetic mesh, which is intersected with a circle to derive a circular mesh. Then, all points of this circular mesh are wrapped onto a sphere. The formation of a curved mesh is an essential step, as it facilitates the wrapping algorithm to identify the points within the mesh closest to the surface of the heart models. After the curved is constructed, the heart model (in this case, a standard tessellation language (STL) file derived from an online repository(*27*)) is placed into the curved auxetic sheet such that they intersect slightly with each other, which improves the wrapping result in the next step. Finally, every point on the auxetic sheet is projected onto the bottom of the heart by computing the point on the heart surface with the smallest distance to a given point on the curved sheet. As with previous shapes, all the different kinds of auxetic meshes can be wrapped onto the heart this way (average design compilation time ∼ 0.2 s) (**Figure 6B**, also see **Figure S4** for additional shape iterations). The corresponding governing equations for the design schemes are provided in the Supplemental Information, and other simpler shapes such as spheres are shown in **Figure S3**. Here, we select the sinusoidal meshes to demonstrate their rapid printability over a heart model (**Figure 6C**). For this, we first volumetrically print a heart using unlabeled GelNB/GelSH resin, followed by projecting supporting pillars and swapping the non-photocrosslinked resin with Rhod-labeled GelNB/GelSH. The auxetic mesh around the heart is then rapidly volumetrically printed **Figure 6D**. Notably, the auxetic meshes in **Figure 6D** were printed directly over the heart to allow better visualization of the mesh by maintaining its shape. We foresee that such complex shapes may one-day pave the way for patient-specific grafts or patches which can conform to the shape of the organ.

**Figure 6.**
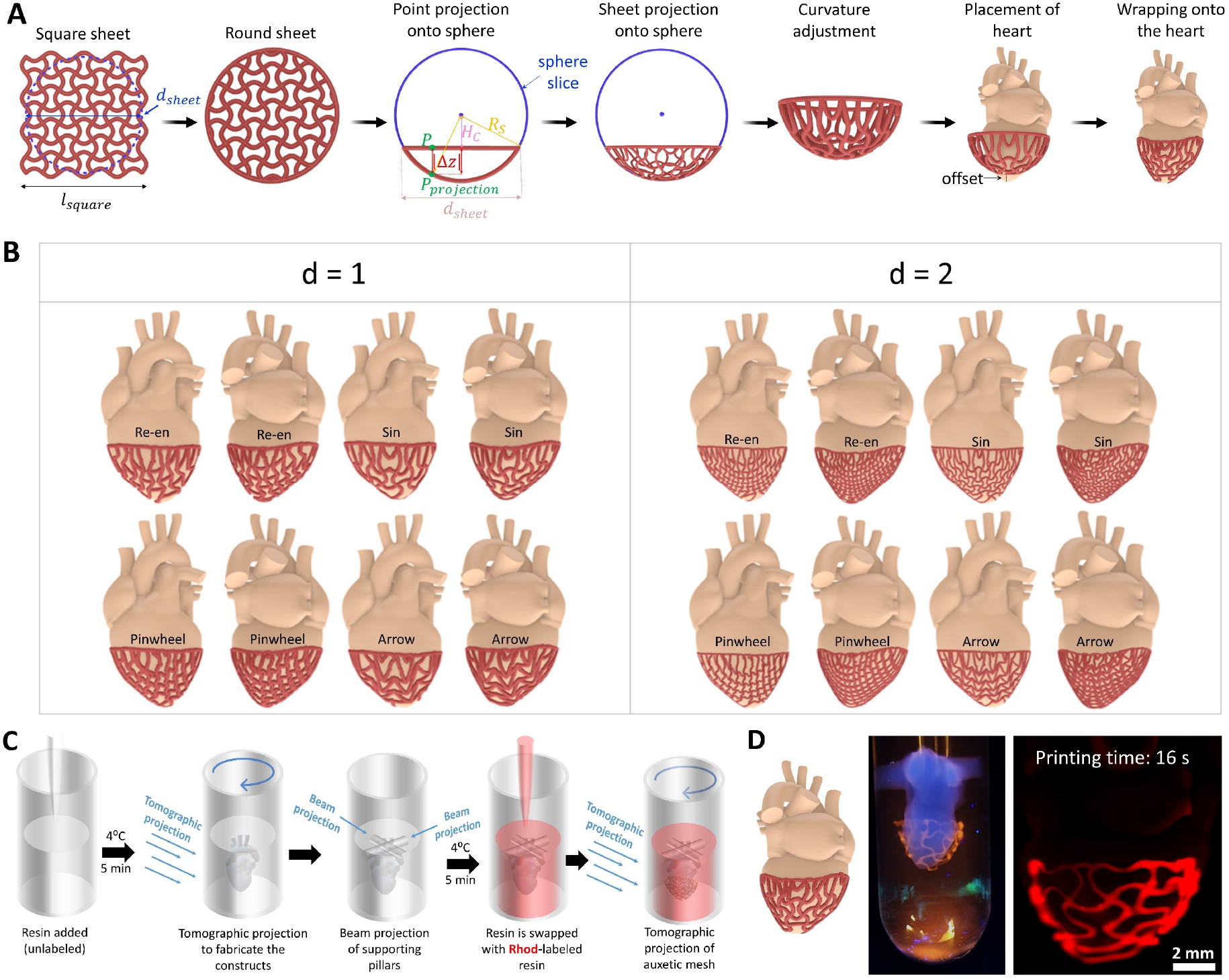
Creation of organ-specific auxetic meshes using wrapping algorithms. **A**. Rationale for the fabrication of the curved auxetic meshes and wrapping the same onto custom organ models (in this case, a heart). **B**. Design iterations of different auxetic meshes wrapped around the heart (d represents the density index of the auxetic structure. **C**. Resin swapping procedure for the fabrication of the heart construct with a wrapped auxetic mesh. **D**. Macroscopic image (left) and light sheet microscopy image (right) of the heart construct (non-labeled) with auxetic mesh (rhod-labeled) around it.

### Algorithmic design of perfusable architectures

Finally, we demonstrate how the algorithmic design and Multimat VP schemes can be used to design and fabricate simple and complex perfusable structures. In the first design scheme, we design a cuboid consisting of a hollow spheroidal center and six perfusable channels around the sphere. The channels are created as a hyperbolic sine function and a circular pattern is created by copying the sine functions 7 times around the center axis (**Figure 7A**). Next, the channels are mirrored with an offset in the middle, followed by connecting the mirrored and original parts through straight channels. By changing the amplitude of the hyperbolic sine function or the diameter of the channels, a variety of perfusable shapes can be created within seconds (**Figure 7B**). These channels are then removed from a cuboidal shape. In addition, a sphere with a pre-set offset with the channels is also removed from the cuboid to result in the perfusable construct with a hollow sphere in-between. To fabricate a construct featuring perfusable channels surrounding a matrix of different material, we introduce the third Multimat VP approach – prefabricated construct integration (**Figure 7C**). In this approach, a FITC-labeled GelNB/GelSH sphere is fabricated, followed by extracting the same and integrating within another resin container partially filled with unlabeled GelNB/GelSH resin. The resin is kept at 24°C to allow easy integration of the sphere. More unlabeled GelNB/GelSH resin is then filled over the construct, and the cuboidal construct with hollow sphere and perfusable channels printed such that the pre-fabricated spherical construct is accommodated within the hollow spherical center in the construct. Perfusing Rhod-labeled GelNB/GelSH into the channels allows us to image the sample using light sheet microscopy, as shown in **Figure 7D**, where the central FITC-labeled sphere is surrounded by a network of Rhod-labeled channels. We foresee that such synergistic algorithmic design and Multimat VP approach can find potential applications in disease-on-a-chip models, for instance, a tumor surrounded by a network of capillaries which demonstrate neovascularization to the tumor site. Such a model can be used to study the effects of biological (macrophages (*28*), exosomes (*29*), etc.) or non-biological therapeutics (*30, 31*) on tumor metastasis or angiogenesis.

**Figure 7.**
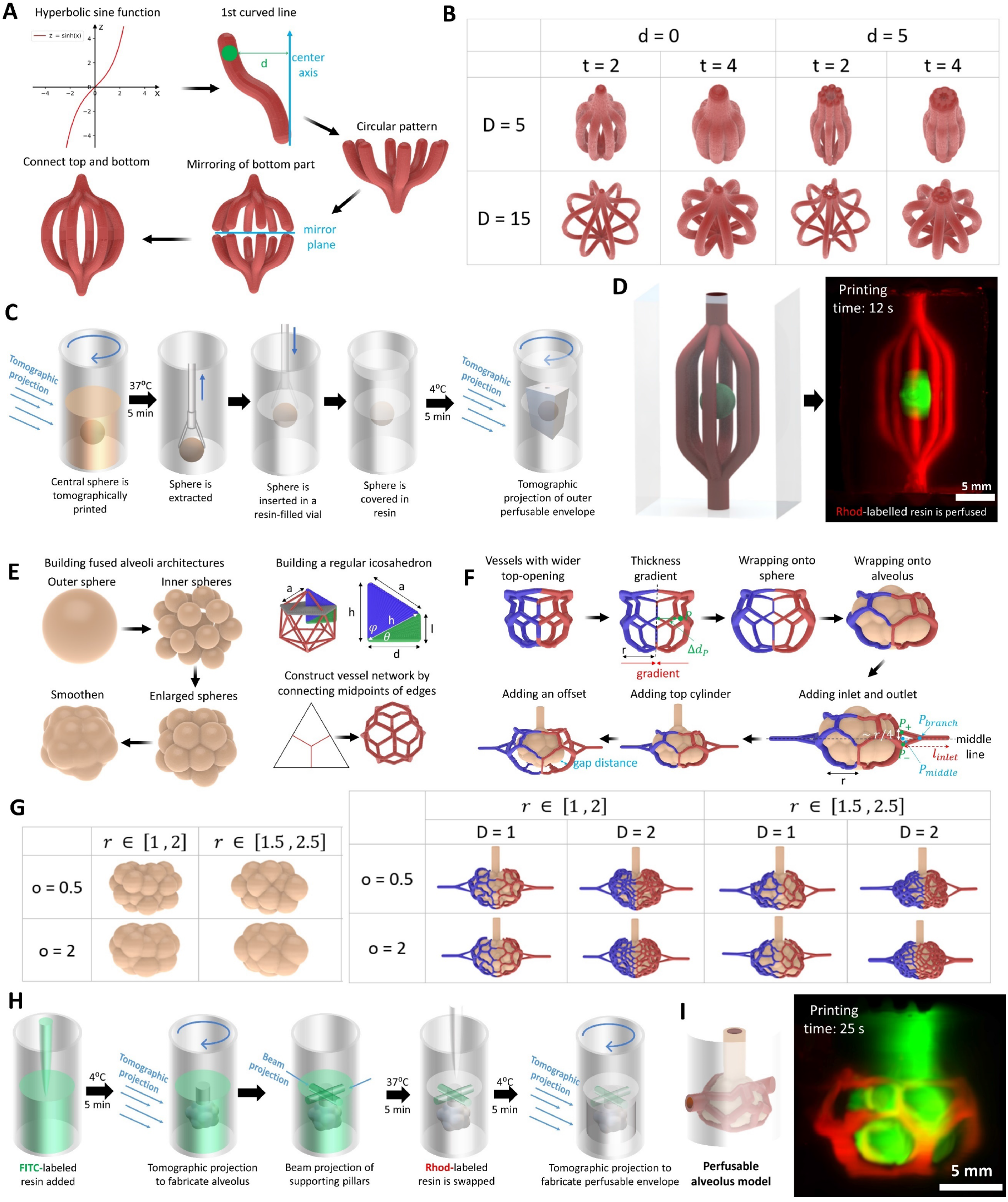
Rapid design and fabrication of perfusable constructs. **A**. Design rationale for simple perfusable architectures featuring multiple bifurcating channels. **B**. Select architectures made under iterations of design parameters (d, t and D represent inlet channel diameter, thickness of individual channels and diameter of the offset of the channels at the center). **C**. Multimat VP scheme, where the central FITC-labeled sphere (GelNB/GelSH) is created first, and then transferred to another resin container pre-filled with unlabeled resin (GelNB/GelSH) and tomographic projections are performed. **D**. Volumetrically printed construct after perfusion with Rhod-labeled resin (images through light sheet microscopy). **E**. Scheme of fabrication of the alveolar budding structures and the perfusable channels. **F**. Wrapping of the perfusable channels around the alveolus. **G**. Shape iterations of alveolar structures (r represents radius of individual mini-spheres and o represents the offset of spheres w.r.t. each other) and the perfusable channels (D represents the density of capillaries) around the alveoli. **H**. Scheme of fabrication of the alveolar construct (also see supplemental **Figure S6**). **I**. Printed construct with FITC-labeled alveolus construct with perfused Rhod-labeled resin surrounding the construct (images captured through light sheet microscopy).

The algorithmic design framework is not just limited to simple perfusable shapes but can also be expanded to more complex shapes such as alveoli surrounded with a perfusable vessel network. The alveolus is created by dividing a sphere into several smaller spheres bound by the periphery of the sphere, followed by scaling-up individual spheres to create the budding alveolus structure (**Figure 7E**). The vessel network surrounding the alveolus is first created as a regular icosahedron (see governing equations in the Supplemental Information). Next, every equilateral triangle along the face of the icosahedron is divided into four identical equilateral triangles by connecting the midpoints of all edges. This process can be repeated to further divide the triangles and create denser structures with smaller hexagons. Then, for every equilateral triangle, its geometric center is connected with the midpoint of each edge to create the vessel network (**Figure 7E**). Further, to improve the quality of prints and avoid intersection with the top cylinder later, the vessel network on the top is enlarged by factor 2 to create a bigger opening (**Figure 7F**). A thickness gradient is introduced to the vessels, such that the thickness decreases gradually from the left and right ends of the shape towards the middle. Then every edge of the icosahedron is interpolated to create 11 equidistant intermediate points, i.e. 10 sub-beams per edge, followed by wrapping each onto the circumscribed sphere of the icosahedron. The result is a spherical vessel shape which can be used for wrapping. The alveolus is placed such that it is concentric with the vessel shape, and every beam on the vessel shape is interpolated into 10 sub-beams and wrapped onto the alveolus by computing the point on the alveolar surface with the smallest distance to a given point on the vessel network. Then a cylinder is created and placed on top of the alveolus, and inlet and outlet ports added such that the inlet/outlet port intersects with a branching point of the vessel network. We can also create an offset by using a larger alveolar shape for wrapping of the vessels, then placing a smaller alveolar shape concentrically for the final shape. By changing the offset, vessel diameters and their densities, we can generate a wide array of alveoli and surrounding vessel shapes (average design compilation time ∼ 0.5 s) as shown in **Figure 7G**. In the model we used for printing, we used a gap of 250 µm between the alveolus and the vessels. The alveoli and the vessel network were removed from a cylindrical construct to create perfusable channels and a hollow portion to accommodate the alveolus. The alveolus construct for printing was also hollowed-out by removing a scaled-down shape from the original alveolus. The printing scheme utilized first printing the alveolar shape, using FITC-labeled GelNB/GelSH, followed by supporting pillar projection (**Figure 7H**). This was followed by swapping resin with unlabeled GelNB/GelSH and printing the hollow construct (see the printed constructs during different stages in **Figure S6**). Finally, Uncrosslinked Rhod-labeled GelNB/GelSH at 37°C is perfused in the channels (**Figure 7I**), and the entire construct exposed to 405 nm UV light to crosslink the resin in the channels. In our future work, we plan to print different vessel densities followed by seeding of epithelial cells to create lung-on-chip models, which can be used to study pulmonary pathologies (*32*).

## Discussion

The algorithmic design scheme offers a transformational approach towards facile creation of a wide array of complex shapes, such as the auxetic meshes and cylinders, organ-specific grafts or perfusable alveoli structures which we demonstrated in this work. The algorithms can rapidly process a wide array of point-point connections (such as wrapping functions, segmentation of icosahedron triangles into smaller triangles, connective lattice elements of auxetic shapes, etc.) which would be tedious and time-consuming to execute manually. Once the design scheme is established for any shape, the shapes can be easily iterated by inputting the ranges and increments for the important parameters, which is otherwise a daunting task to perform in conventional CAD softwares. This difference in designing and iterating is even more profound as the shape complexity increases, with the introduction of organ-specific auxetic meshes and the alveolar structures. Here the algorithms ensure that the interconnections between the constitutive points or contours are satisfied when a new shape iteration is formed. We have shown that algorithm-based computational modeling schemes can allow rapid screening of the mechanical properties of large arrays of complex architectures. Here, integration of the iterative algorithmic design and computational modeling within a deep learning framework can lead to a powerful framework for shape optimization(*33*). Our future work will entail expanding the computational modeling to simulate fluid flow within perfusable architectures.

We have made the design and simulation code available in the GitHub repository (see “**Data and materials availability**” section). For users who are not adept at coding, we have created graphical user interfaces where the users can load their own models, to be able to design their own custom auxetic patches or perfusable networks with varying vessel densities. Using the interface, the users can also iterate the designs of the auxetic and perfusable shapes demonstrated in the present work for their own applications. In addition, dedicated libraries have been established which can be used for executing the structural simulations or for creating porosities within constructs (see **Figure S5**).

In this work, we deployed three techniques for the rapid volumetric printing of multi-material constructs: **1**. Tomographic projection printing of first layer of the construct supported by projected pillars, followed by swapping of the uncrosslinked resin with a different composition and tomographic projection printing of the subsequent layer. **2**. Filling the printing vials with two different resin compositions and tomographic projection printing of the entire construct at once. **3**. Incorporating a prefabricated construct into a resin-filled container, followed by tomographic projections to fabricate the multimaterial constructs. The rationale for which Multimat VP method needs to be used would depend on the complexity of the structure that needs to be fabricated, and the resin composition. While the first scheme offers tremendous design freedom (we printed multi-material auxetic cylinders (Figure 4) auxetic mesh on heart (Figure 5) or perfusable alveolar models (Figure 6) using this method), aligning subsequent projections with the first projection is often challenging. Furthermore, removal of the supporting beams after printing can lead to material losses and may even be difficult to execute for fragile constructs. For multi-material constructs such as tissue interfaces, the second scheme can be a more robust scheme to align the two layers (such as the bilayered auxetic meshes in Figure 3). However, for resin compositions which do not undergo thermo-reversible crosslinking, the second scheme may cause mixing of the two resins across different regions, especially when the resin viscosity is low. In such a case, the remaining two strategies are a better choice. Notably, we have used the same base material of the resin when performing more than one tomographic projection to make the constructs (i.e., first and third strategy). Since the change in RI before and after crosslinking of GelNB/GelSH resin was small (ΔRI = 0.002), we did not observe a substantial difference in resolution between the first and second projections. However, if the base material is different between the projections, the differences in RI between the photocrosslinked constructs can cause unwanted scattering or diffraction of light which may affect the print resolution and quality. Similar constraints may also apply to the third strategy of prefabricated construct integration. In this case, RI matching agents such as Iodixanol (*19*) can be used to fine-tune the RI of the resins and achieve high resolution prints.

While we only demonstrated two resin compositions within the printed constructs in the present work, the techniques can be easily adapted to more than two material compositions to create more complex constructs. For example, bioinks pertaining to bone, tendon and muscle can be sequentially added into the print vial for the fabrication of bone-tendon-muscle interfaces. Here, while the bone construct could feature a porous matrix (see foaming algorithm outcomes in **Figure S5**), a fascicular arrangement of muscle and tendon may be difficult to achieve via tomographic projections. Therein, hybridization of VP with filamented light projection (FLight, a technology developed in our group (*34*)) could allow one to fabricate the muscle and tendon interfaces with fascicular arrangement of muscle fibers or aligned collagen in tendons, while the bone can be tomographically projected to create a porous matrix. In fact, hybridization can also be performed with other printing techniques (*35, 36*). For example, extrusion printing of photoresins can be used to control the spatial distribution of different materials within the resin container. Subsequently, single tomographic projection can be used to create the multimaterial constructs. Such hybridization schemes will be the future scope of our investigation. Naturally, for tissue engineering purposes, imparting a macroscopic vasculature and further inducing neovascularization (*10, 37*) will be a key aspect to allow physiological-scale tissue fabrication.

For this work, in order to demonstrate rapid printing, photoclick materials based on step-growth polymerization were ideal as we have established expertise on high speed volumetric printing using these materials (*17*). However, the materials demonstrated in this work may not be suitable towards all biomedical or structural applications as the modulus is small (∼ 10-100 kPa). Herein, one could also use chain-growth polymerization, which is typically found in acrylate or methacrylate-based resins, to obtain stiffer structures (*38*) with minor compromises on the fabrication duration as the rate of photopolymerization is slow (e.g. volumetric printing of gelatin methacrylate takes ∼ 30 s/cm^3^ of construct, while GelNB/GelSH take ∼ 8 s/cm^3^ of constructs (*17*)). Future research on photoclick-compatible materials or their hybridization schemes, which yield higher stiffness constructs, could improve the applicability of the materials to a wider variety of biomedical applications such as polymer-based arterial stents (*39*) or tracheal grafts (*40*). Further, current volumetric printing approaches have been limited to constructs spanning only a few centimeters in sizes, and future research on tomographic projections within larger containers can circumvent such size limitations. As such, one also does not need to use volumetric printing or deploy photocrosslinkable materials. The shapes generated and optimized through the algorithmic design and computational modeling schemes can be fabricated though conventional or bespoke manufacturing processes integrated into larger assembly lines, as long as the complexity of the shape can be achieved. For example, the algorithmic schemes could help speed up product design and optimization of prosthetic implants or metallic stents, to even vehicle drive shafts and engines, which could bring about substantial cost-savings in the product pipeline. This work can potentially transform the way engineers or scientists approach new design problems and develop solutions that have the potential to benefit society at large.

## Materials and Methods

### Algorithmic design

Hyperganic core (an algorithmic engineering platform developed by Hyperganic GmbH) environment was used to run the algorithmic design schemes written in C#. Detailed explanations and equations of the algorithms are provided in the Supplemental Information. We have created graphical user interfaces within Hyperganic core which will allow users to change the designs of the auxetic and perfusable shapes. User Interfaces have been created for each design architecture with provision to create designs based on variations of design parameters (h, t, v, n1, n2, etc.), and are integrated in the Hyperganic source code shaped on the open-source library. See “Data and materials availability” section for the sources codes for the design schemes and procedures for opening the graphical user interfaces to change different designs.

### Computational modeling of the structural mechanics of the auxetic meshes

Numerical analysis of auxetic meshes has been performed using the simulation kernel of Hyperganic Core (governing equations for the models are provided in the **Supplementary Information**). The simulation functionality is integrated within the C# API that directly integrates with the algorithmic design schemes. See “Data and materials availability” section for the sources codes and procedures for running the codes.

### Matrix synthesis

The norbornene or thiol-modified gelatin were synthesized using procedures previously established in our lab (*17, 41*). For formulation of GelNB, porcine-derived (Type A) gelatin was dissolved in 0.5 M carbonate-bicarbonate buffer (pH∼9, obtained by adding 38.2 g/l of sodium bicarbonate and 4.7d/l of sodium carbonate in deionized (DI) water) at 50°C to get a 10% w/v solution. After obtaining a clear solution under stirring, carbic anhydride was added at a concentration of 100 mg/g of gelatin. After letting the reaction proceed for 1 h, the solution was dialyzed (at 40°C) with frequent DI water changes (2 per day) for 5 days. The matrix was then lyophilized for 4 days and stored at -20°C until further use. For formulation of GelSH, porcine-derived (Type A) gelatin was dissolved in 0.15 M MES (2-(N-morpholino)ethanesulfonic acid) buffer (pH∼4) at 50°C to get a 2% w/v solution. When completely dissolved, DTPHY (3,3’-Dithiobis(propionohydrazide)) was added at 95 mg/g of gelatin while stirring. When completely dissolved, EDC (1-Ethyl-3-(3-dimethylaminopropyl)carbodiimide) was added at 135 mg/g of gelatin while stirring. The reaction is then allow to proceed at 50°C under stirring for 12 h. Next, TCEP was added at 344 mg/g of gelatin, followed by continuing the reduction reaction for 6 hours. Finally, 1g of NaCl was added and the solution dialysed against DI water balanced to pH 4.5 with diluted HCl. The degree of functionalization of the matrices was determined using ^1^H NMR spectroscopy (GelSH DS: 0.276 ± 0.016 mmol/g, GelNB DS: 0.217 ± 0.007 mmol/g, plots provided in supplemental information **Figure S7**).

The GelNB/GelSH resin was formulated by mixing the lyophilized GelNB or GelSH matrix in PBS to achieve 5% w/v total gelatin concentration. GelNB/PEGSH resin was formulated by mixing the lyophilized GelNB matrix in PBS at 3.8% w/v and thiolated 4-arm PEG (10 kDa, SinoPEG) at 1.2% w/v. For both resin formulations, 0.05% w/v LAP (Lithium phenyl(2,4,6-trimethylbenzoyl)phosphinate) was used as the photoinitiator. Rhod- or FITC-labeled resin was formulated by adding Acryloxyethyl thiocarbamoyl Rhodamine B (Rhod-Acr) or Fluorescein isothiocyanate (FITC) stock in DMSO (at 10 mg/ml) to the resin formulation at 1 µl/ml. FITC is conjugated to the resin matrix through amide bond formation with the primary amines of the matrix, while Rhod-Acr conjugates to GelSH through thiol-ene reaction during printing. The amounts of FITC and Rhod-Acr do not affect the light dose of the resin compared to a non-labeled resin.

### Multimat volumetric printing

The open format volumetric printer from Readily3D was used for these experiments. This printer allowed for each resin swapping and visualization of the constructs during and after printing. Procedures for Multimat VP have been discussed in the results section. Here we provide necessary details for replicability. Prior to printing, dose tests were performed by projecting an array of circles (ɸ = 1 mm) featuring a variation of light intensity onto a 3 mm path length cuvette filled with the resin. Diameters of the projected cylinders were then measured using bright field microscopy and the light dose allowing for the diameter to be 1 ± 0.05 mm (dimensions measured in ImageJ) was chosen. For all prints, 18 mm printing vials were used. After filling-in the desired volume of the resin – 4 ml per layer for single material VPs and 3 ml per material for the MultiMat VP experiments, the resin was allowed to thermo-reversibly crosslink for 5 min at 4°C. This allowed stability of the printed structure during photocrosslinking. To remove the non-crosslinked resin (to swap the resin in between different tomographic projections or at the end of the printing), the vial was kept at 37°C for 5 min, followed by washing with warm (37°C) PBS twice. The supporting pillars were fabricated in a sequential manner: First a single circular beam projection at highest laser power permissible by the printing system (64 mW/cm^2^) was used for 10 s to print the first pillar. The vial was then rotated by 90° and the projection performed again.

### Cell culture and tissue immunohistochemistry

C2C12 murine myoblasts and NIH 3T3 murine fibroblasts were cultured in Dulbecco’s Modified Eagle Medium (DMEM) supplemented with 10% w/v fetal bovine serum (FBS) and 1% w/v penicillin/streptomycin. The cells were passaged at 80% confluency using 0.25% w/v trypsin and 0.05% w/v EDTA. Next, C2C12 and 3T3 cells were labeled with CellTracker™ red and green dyes (ThermoFisher), respectively, using manufacturer’s specifications. The cells were then resuspended in GelNB/GelSH matrix at 2.5 M cells/ml and the muscle-connective tissue units volumetrically printed. After 4 weeks of culture, muscle-connective tissue units were immunohistochemically stained for myosin heavy chain (MyoHC), Collagen I, Actin filament (Phalloidin) and nuclei (DAPI) based on our previous work (*34*).

### Light Sheet Microscopy

An axially scanned light sheet microscope (MesoSPIM, V4)(*42*) was used to image fluorescently labelled samples. The constructs were mounted onto a custom printed microscope sample holder and submerged in a quartz cuvette filled with mQ water, which was then mounted onto the MesoSPIM microscope stand. For imaging, a macro-zoom system (Olympus MVX-10) and 1x air objective (Olympus MVPLAPO1x) with adjustable zoom were used. Voltage adjustments using the electrically tunable lens (ETL) were performed for each run. Step size was chosen from 10 – 50 µm.

## Acknowledgments

P.C. acknowledges a Marie Skłodowska Curie postdoctoral fellowship (grant number 101024341). M.Z.W. acknowledges ETH Grant application ETH-38 19-1 and Innosuisse funding application no. 55019.1 IP-ENG for their kind support. D.R. acknowledges Swiss National Science Foundation project grant 205321_179012. The authors further acknowledge the assistance from ETH (ScopeM) imaging facility, and UZH MesoSPIM light sheet microscopy initiative. We thank Hyperganic Group GmbH for providing the free academic license to use their Hyperganic Core software.

## Author contributions

Conceptualization: P.C., M.Z.W.

Methodology: P.C., D.R., A.W., R.R., H.L., P.W., D.D., N.K., M.Z.W.

Investigation: P.C., D.R., A.W., R.R., H.L., P.W., N.K.

Visualization: P.C., D.R., A.W., N.K.

Supervision: M.Z.W., P.C., N.K.

Writing—original draft: P.C., D.R., A.W.

Writing—review & editing: M.Z.W., P.C.

## Data and materials availability

All data are available in the main text or the supplementary materials. The source code for the designs and simulations and their graphical user interfaces, as well as instructions on how to run the code and GUIs have been provided in the GitHub repository: https://gitlab.hyperganic.com/hyperganic-education/hyperganic-partners/auxetic_and_perfusable_shapes. Academic users can contact Hyperganic to obtain login and software access. Other data is available on this online repository: https://www.research-collection.ethz.ch/handle/20.500.11850/583621 (DOI: 10.3929/ethz-b-000583621).

