## Supplemental information for "Synergizing algorithmic design, photoclick chemistry and multi-material volumetric printing for accelerating complex shape engineering"

### Supplementary Materials for

#### Synergizing algorithmic design, photoclick chemistry and multi-material volumetric printing for the rapid fabrication of complex structures

Parth Chansoria[1], Dominic Rütsche[1,2], Anny Wang[1,3], Hao Liu[1], Davide D'Angella[3], Riccardo Rizzo[1], Amelia Hasenauer[1], Patrick Weber[1], Nafeesah Ibrahim[1], Nina Korshunova[3], Marcy Zenobi-Wong[1*]

**This PDF file includes:**
Figures S1 to S7.

Governing equations for the algorithmic design schemes and computational models to determine the Poisson's ratios.

The code and graphical user interfaces for building the different auxetic and perfusable can be found in the GitHub repository: https://gitlab.hyperganic.com/hyperganic-education/hyperganic-partners/auxetic_and_perfusable_shapes. Academic users may contact Hyperganic to obtain the login credentials and software access.

.

### Supplementary Figures

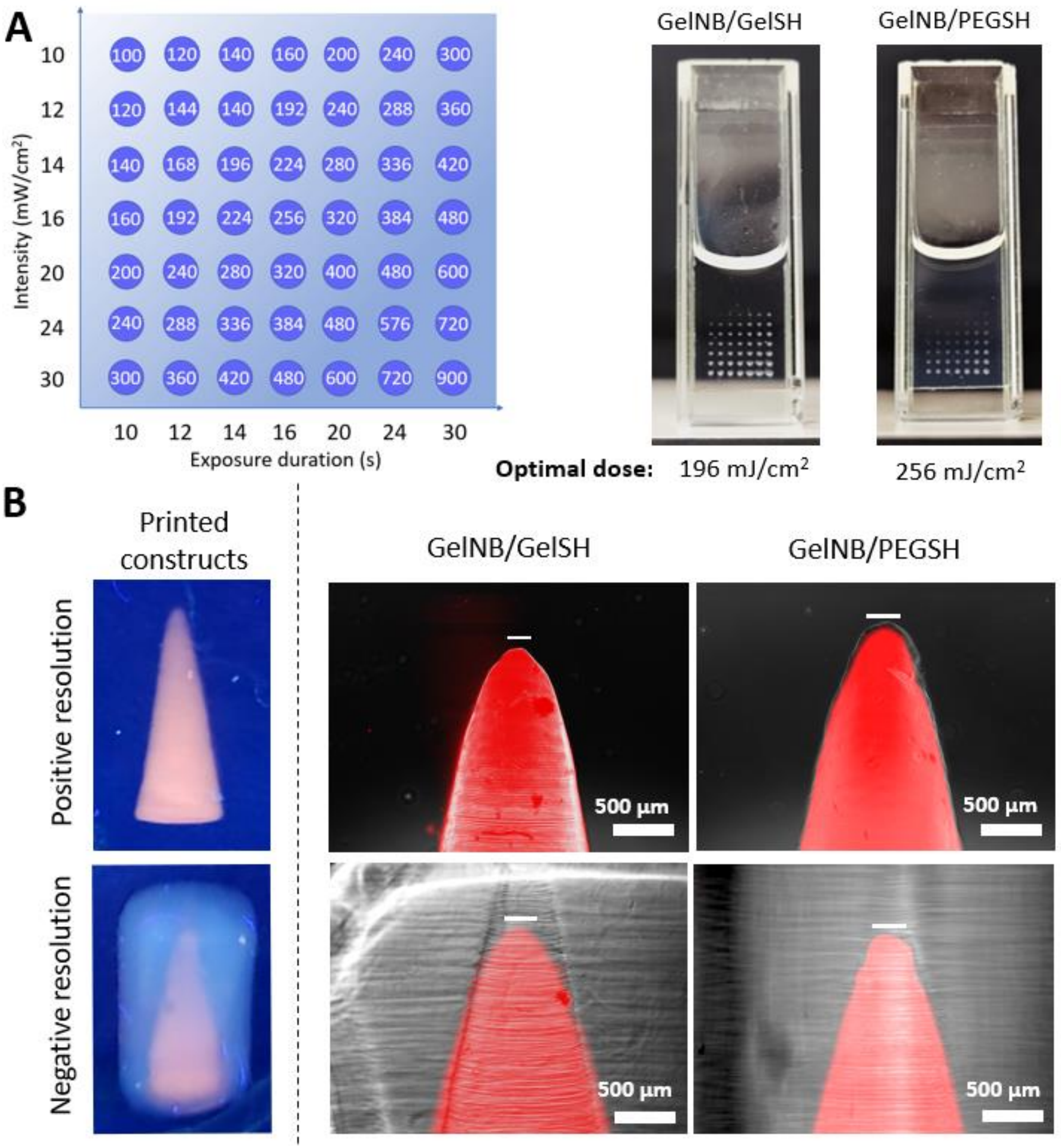


**Figure S1. A.** Dose tests for the different photoclick resins, and the identified optimal light doses. **B.** Positive and negative resolution tests demonstrate that upto 125 µm positive resolution and 200 µm negative resolution is achievable with the GelNB/GelSH resin, while upto 200 µm positive and negative resolution is achievable with the GelNB/PEGSH resin.

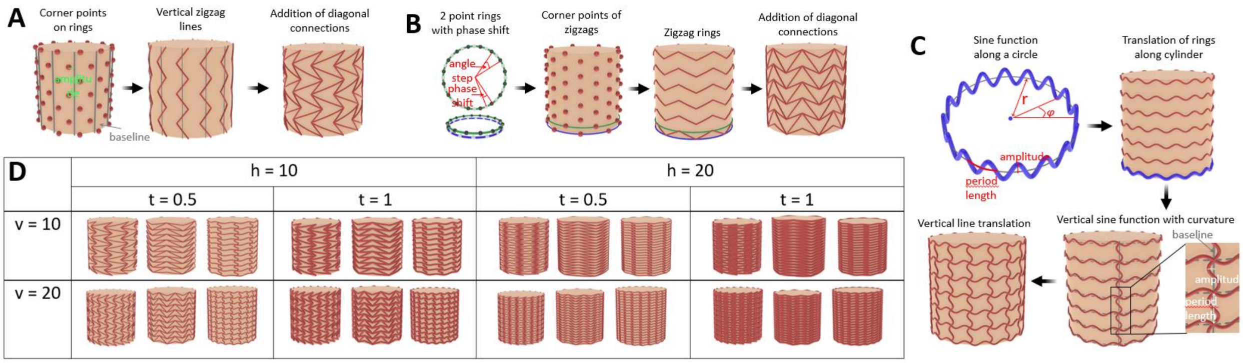


**Figure S2.** Algorithmic design scheme for cylinders featuring: **A.** horizontally oriented arrowhead elements, **B.** vertically oriented arrowhead elements, and **C.** pinwheel elements. **D.** A variety of shapes can be generated within seconds by varying the thickness (t) of the elements, number of vertical lines (v) in a ring and number of horizontal lines (h) from the top to the bottom.

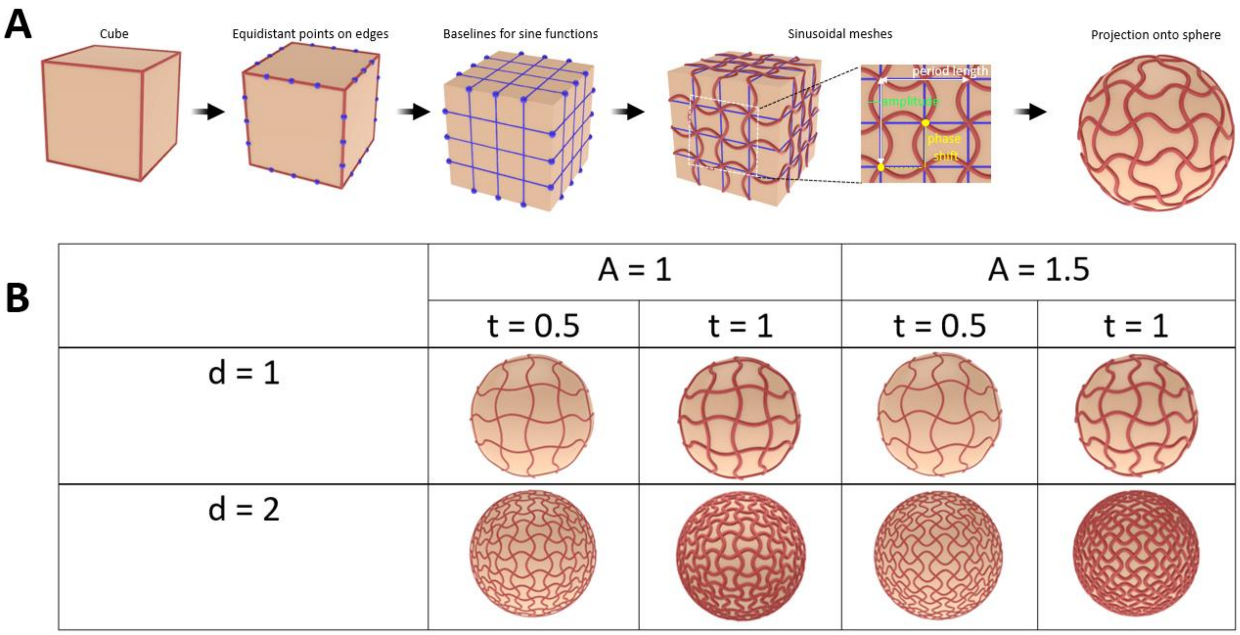


**Figure S3. A.** Algorithmic design scheme for creating auxetic meshes around spherical shapes, and their design iterations. **B.** A variety of shapes can be generated within seconds by varying the amplitude (A) amplitude of sinusoidal lines, line thickness (t), and density index (d) of the auxetic structure (relative, indicates the level of density).

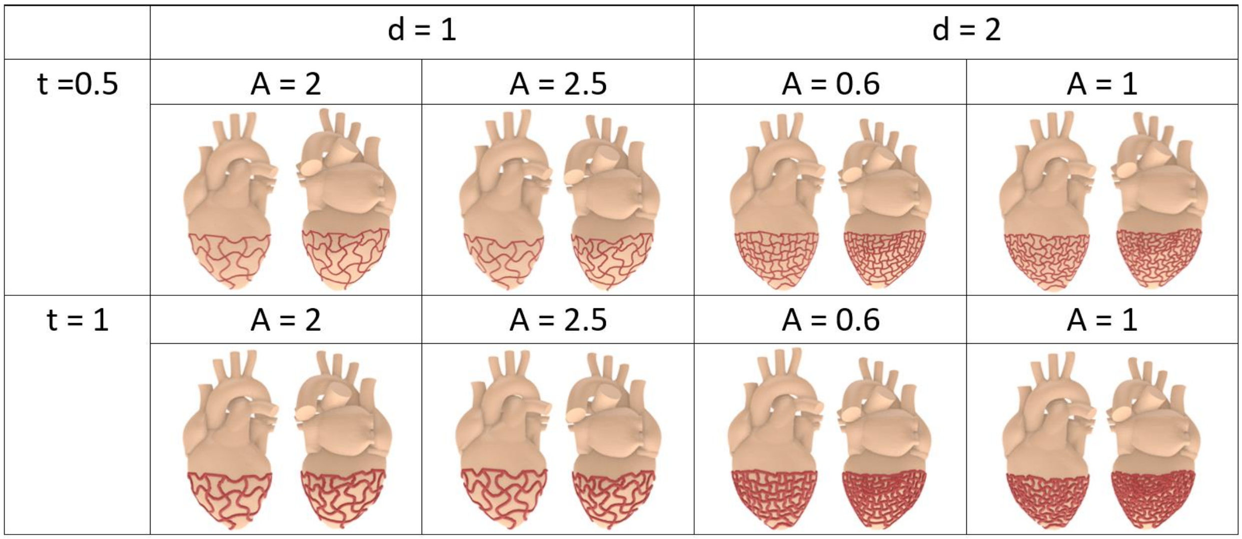


**Figure S4.** The sphere can be replaced with the heart model and the wrapping algorithm (see methods in the Subsection "Design rationales for different shapes") can be used to wrap the designs around the heart model. A variety of shapes can then be generated by changing the amplitude of sinusoidal lines (A), line thickness (t), and density index (d) of the auxetic structure (relative, indicates the level of density).

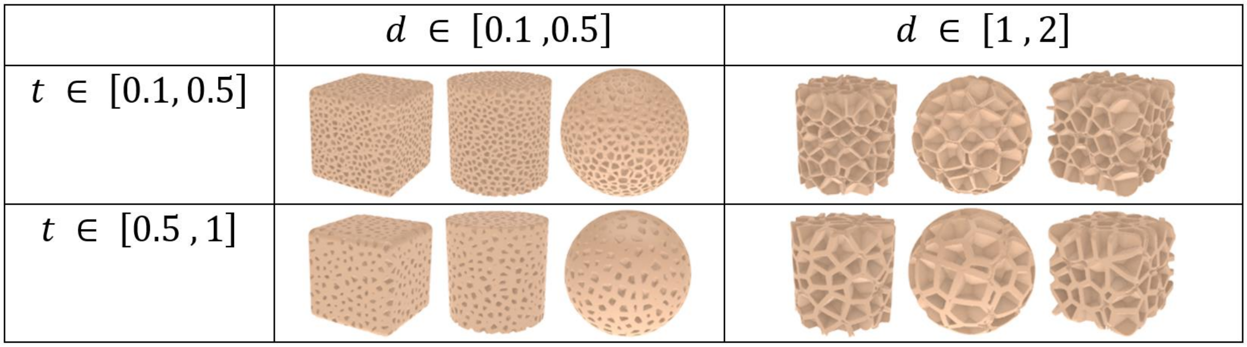


**Figure S5.** Example shapes generated using the foaming algorithm (see methods in the Subsection "Design rationales for different shapes"). The varied parameters include distance between cell centers (d) and the wall thickness (t). See supplemental section "Foamed Shape" for additional information on library to be used.

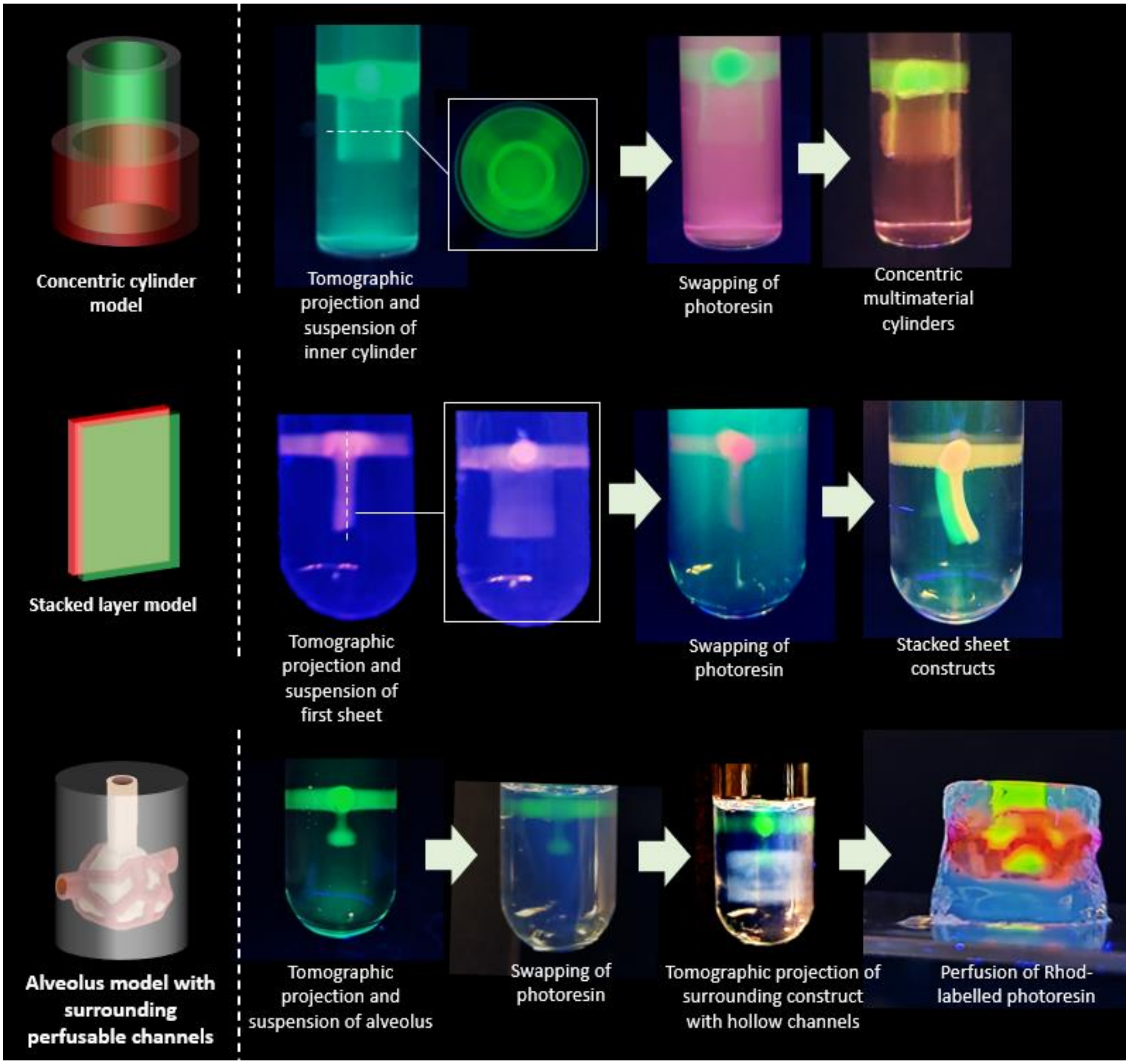


**Figure S6.** Example constructs fabricated using the resin swapping scheme which allows the material and the design of the printed constructs to be changed across their thickness.

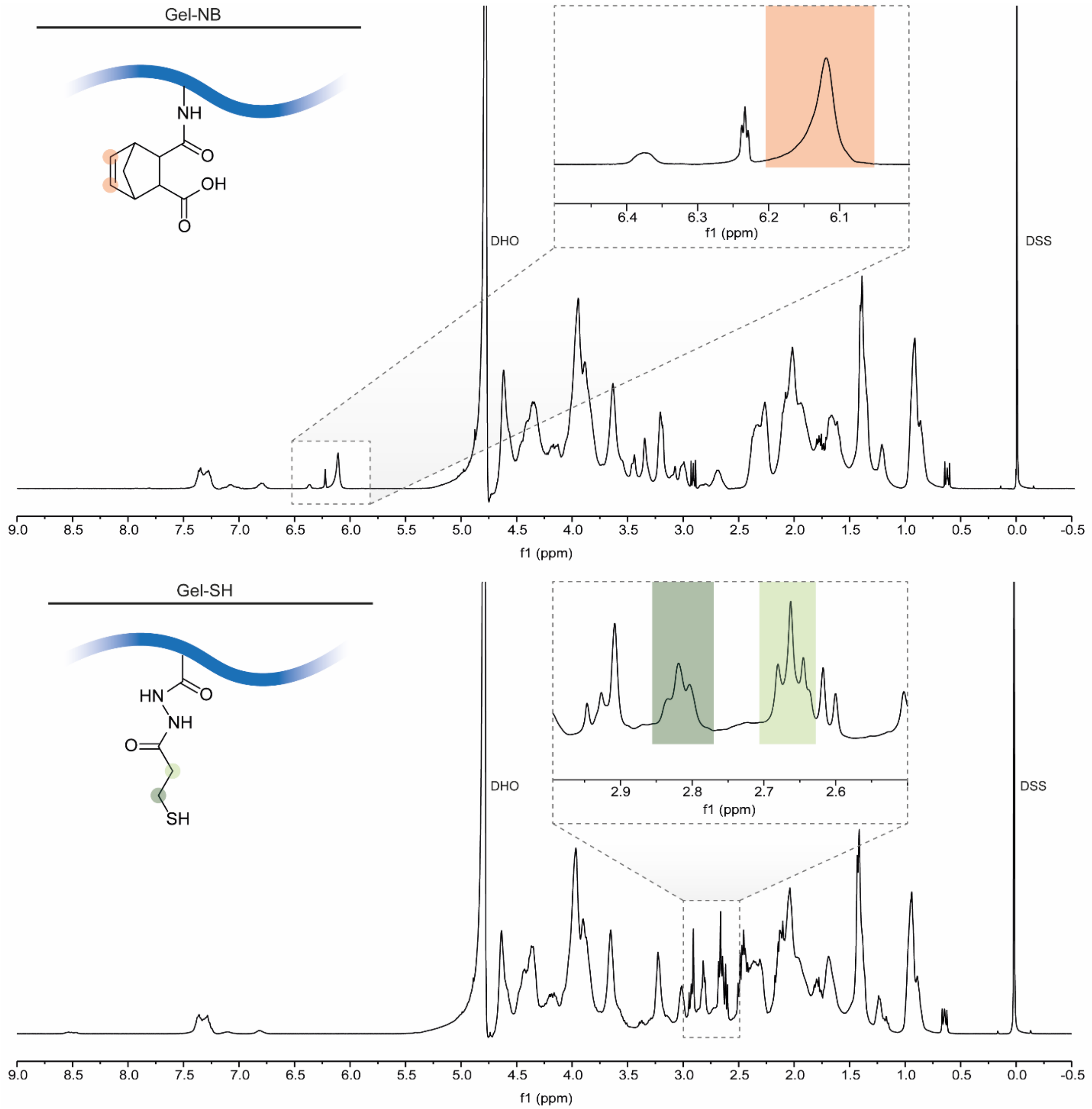


**Figure S7.** $^{1}H$ NMR spectra of GelNB and GelSH. For GelNB, the integral of NB-ene protons (orange, close up) was compared to the integral of DSS internal standard methyl protons (0.1 to -0.1 ppm) to calculate the degree of substitution (DS), as per our previous work (*1*). For GelSH, the DS was calculated by comparing the integrals of the DTPHY methylene protons (green, close up) the integral of DSS internal standard methyl protons (0.1 to -0.1 ppm), as per our previous work (*2*).

### Design rationales for different shapes

***Auxetic mesh: Re-entrant honeycomb***

The auxetic sheet is constructed on xy-plane, with all points having $z = 0$. The structure can be interpreted as vertical zigzag lines in z-direction connected by horizontal beams in x-direction. All points on a zigzag line can be created by translation of equidistant points on the baseline.

First, points on the baselines of the zigzags in y-direction are created. Given the y-dimension of the sheet and the number of steps along x-direction, the vertical distance between two neighboring points on a baseline is determined by

$$vertical\ distance = \frac{y - dimension\ of\ sheet}{step\ number\ in\ y - direction}$$

Given the x-dimension of the sheet and the number of zigzag lines, the distance between two neighboring baselines is

$$horizontal\ distance = \frac{x - dimension\ of\ sheet}{number\ of\ zigzag\ lines}$$

After defining all points on the baselines, the zigzag lines can be created by translating all points along x-direction with an amplitude of

$$amplitude = \ horizontal\ distance/4$$

Two consecutive points on a baseline are translated with a positive and negative amplitude respectively to obtain the zigzag structure. Furthermore, all points on two neighboring zigzag lines undergo the opposite amplitude. Hence, for every point with a positive amplitude translation on the baselines, all its neighboring point in x- and y- directions undergo a negative amplitude translation, vice versa. After the translation, all points from the same baseline are connected to construct the vertical zigzag lines. Finally, all points on a zigzag line are connected to one of their neighbors in x-direction in an alternating fashion to obtain the final shape.

**Auxetic mesh: Arrowhead**

The auxetic sheet is constructed on xy-plane, all points will have $z = 0$. The structure can be interpreted as vertical zigzag lines in z-direction connected by diagonal beams. All points on a zigzag line can be created by translation of equidistant points on the baseline.

First, points on the baselines of the zigzags in y-direction are created. Given the y-dimension of the sheet and the number of steps along x-direction, the vertical distance between two neighboring points on a baseline is determined by

$$vertical\ distance = \frac{y - dimension\ of\ sheet}{step\ number\ in\ y - direction}$$

Given the x-dimension of the sheet and the number of zigzag lines, the distance between two neighboring baselines is

$$horizontal\ distance = \frac{x - dimension\ of\ sheet}{number\ of\ zigzag\ lines}$$

After defining all points on the baselines, the zigzag lines can be created by translating all points along x-direction with an amplitude of

$$amplitude = \ horizontal\ distance/4$$

Two consecutive points on a baseline are translated with a positive and negative amplitude respectively to obtain the zigzag structure. After the translation, all points from the same baseline are connected to construct the vertical zigzag lines. Finally, every second point on a zigzag line is connected to two points on the zigzag line in positive x-direction.

**Auxetic mesh: Sinusoidal ligaments**
The sinusoidal ligaments consist of sine functions in x- and y-direction. All points will have $z = 0$. The crossing points of the functions correspond to the crossing points of their baselines.

First, the baseline positions of the sinusoidal lines in x-direction are defined. The vertical distance between the lines is

$$vertical\ distance = \frac{y - dimension\ of\ sheet}{step\ number\ in\ y - direction}$$

The chosen amplitude of the sine function is

$$amplitude = \ vertical\ distance/4$$

Then the coordinates of all points on one sinusoidal line are

$$x\ \in [0, x - dimension\ of\ sheet]$$

$$y = 0$$

$$y = amplitude * \sin(period * x\ + phase\ shift) + y_{baseline}$$

$$period = \frac{2 * \pi}{period\ length}$$

$$period\ length = 2 * vertical\ distance$$

The $phase\ shift$ is 0 and $\pi$ for two neighboring lines to match the minima of one line with the maxima of the other. All sinusoidal lines in z direction can be constructed following the same principle above, by simply exchanging all x- and y-related variables and replacing vertical distances with horizontal distances.

***Auxetic mesh: Pinwheel***

The sinusoidal ligaments consist of sine functions in x- and y-direction. All points will have $z = 0$. The crossing points of the functions correspond to the crossing points of their baselines.

First, the baseline positions of the sinusoidal lines in x-direction are defined. The vertical distance between the lines is

$$vertical\ distance = \frac{y - dimension\ of\ sheet}{step\ number\ in\ y - direction}$$

The chosen amplitude of the sine function is

$$amplitude = \ vertical\ distance/8$$

Then the coordinates of all points on one sinusoidal line are

$$x\ \in [0, x - dimension\ of\ sheet]$$

$$y = 0$$

$$y = amplitude * \sin(period * x) + y_{baseline}$$

$$period = \frac{2 * \pi}{period\ length}$$

$$period\ length = vertical\ distance$$

All sinusoidal lines in y-direction can be constructed following the same principle above, by simply exchanging all x- and y-related variables and replacing vertical distances with horizontal distances.

***Auxetic cylinders with vertically oriented re-entrant honeycomb elements***

All cylinders mentioned below are aligned along and centered around the z-axis. In the following computations, the conversion from cartesian coordinate system to cylindrical coordinate system to describe points on a cylinder with radius $r$ is:

$$x = r * \ \cos(\varphi)$$
$$y = r * \ \sin(\varphi)$$
$$z = z$$

On a given plane, the azimuth $\varphi$ is the angle between the reference direction $(x, 0, 0)$ and the line from $(0, 0, z)$ to $(x, y, z)$.

First, create the vertical zigzag lines by defining the coordinates of all corner points. The corner points of every zigzag line can be described as points alternating on two sides of a base line. Hence, the x- and y coordinates of corner points are defined by

$$x = r * \cos(\varphi_{baseline} \pm amplitude)$$

$$y = r * \sin(\varphi_{baseline} \pm amplitude)$$

$Amplitude$ represents the distance between a corner point and the baseline in terms of angle, its sign alternates along the z-direction. The corners of the zigzag lines lie on circles parallel to the xy-plane. The x- and y-coordinates of all points on a circle can be obtained by iterating around the circle with a certain angle step length. The total number of iterations (n) necessary to create all points on one ring is:

$$n = \frac{2 * \pi}{angle\ step}$$

$Angle\ step \in [0, 2\pi]$ is the difference in angle between two neighboring baselines. To create the symmetry of the zigzag structure, $angle\ step$ is set as

$$angle\ step = 2 * amplitude$$

For $i \in [0,\ n-1]$, the x- and y-coordinates of all baselines are

$$x_{baseline} = r * \cos(i * angle\ step)$$

$$y_{baseline} = r * \sin(i * angle\ step)$$

Therefore, the x- and y-coordinates of all corner points on the bottom circle are

$$x = r * \cos(i * angle\ step \pm amplitude)$$

$$y = r * \sin(i * angle\ step \pm amplitude)$$

The second circle from the bottom has a certain phase shift to the first one, which can be introduced by the following coordinate formulas:

$$x = r * \cos(i * angle\ step \mp amplitude)$$

$$y = r * \sin(i * angle\ step \mp amplitude)$$

i.e., the sign of amplitude is opposite for two points with the same index in the first and second circle. Since the only constantly changing coordinate is the z-coordinate, all circles can be created by translating the first two along the z-axis. After all the corner points of the vertical zigzag lines are defined, these are connected with straight beams. To match the cylinder curvature, every point on the straight initial beam is wrapped onto the cylinder coat individually using the following formula:

$$p_{wrapping} = p_{initial} + r * \vec{e}$$

Where $p_{wrapping}$ and $p_{initial}$ are on the same plane parallel to xy-plane, and $\vec{e}$ is the unit vector with direction from the circle center $p_{center}$ to $p_{initial}$, and can be computed from vector $\vec{a}$:

$$\vec{a} = p_{initial} - p_{center}$$

$$\vec{e} = \frac{\vec{a}}{|\vec{a}|}$$

All vertical zigzag lines are generated by connecting the corner points using this method. Finally, to create the horizontal beams, every second pair of neighboring points on each ring of the cylinder is connected by an arc using the wrapping method above. Moreover, the position of these horizontal beams alternate along the z-direction to eventually create the re-entrant honeycomb shape.

***Auxetic cylinders with horizontally oriented re-entrant honeycomb elements***

First, create zigzag rings around the cylinder by determining coordinates of the corner points. The corner points of each zigzag ring are composed of two circles of points with a certain phase shift to each other. The total number of iterations necessary to create all points on one circle is:

$$n = \frac{2 * \pi}{angle\ step}$$

$Angle\ step$ is the difference in angle between two neighboring points in a circle. For $i \in [0,\ n-1]$, the x- and y-coordinates of the corner points on the first circle can be calculated by

$$x = r * \cos(i * angle\ step)$$

$$y = r * \cos(i * angle\ step)$$

Whereas the corner points on the second circle have the coordinates

$$x = r * \cos(i * angle\ step + phase\ shift)$$

$$y = r * \cos(i * angle\ step + phase\ shift)$$

To create a re-entrant honeycomb structure with horizontally oriented elements, the phase shift is set to

$$phase\ shift = \frac{angle\ step}{2}$$

Then, one can translate coordinates from the first two circles vertically to create all circles along the z-direction. For this honeycomb structure, the points on two consecutive circles have the same x- and y-coordinates, while two pairs of circles on either side of the above circles have a phase shift. After defining all corner points of the zigzag rings, these can be connected to create the zigzag rings. Before generating the connecting beams, every beam between two corner points is wrapped onto the cylinder coat using the wrapping method mentioned in re-entrant honeycomb cylinder with vertically oriented elements. Finally, vertical beams are constructed between neighboring zigzag rings, i.e., for all corner points in a zigzag ring, there are alternating beams in positive and negative z-directions to the neighboring rings.

***Auxetic cylinders with sinusoidal ligament elements***

First, create a circle parallel to xy-plane. The x- and y-coordinates of all points on the circle vary with the angle $\varphi$, and the z-coordinates correspond to a sinusoidal function along the circumference. For a circle with radius $r$, the coordinates of each point on the circle are

$$x = r * \cos(\varphi)$$

$$y = r * \sin(\varphi)$$

$$\mathrm{z} = amplitude * \sin(period * \varphi)$$

$$period = \frac{2\pi}{period\ length}$$

where $\varphi \in [0,\ 2\pi]$, $period\ length \in [0, 2\pi r]$. Then a second circle with a certain phase shift to the first one is generated, such that the minima of the second circle are exactly over the maxima of the first one. While the formulas for the x- and y-coordinates on the second circle are the same as above, the z-coordinates can be calculated with the following formula:

$$z = amplitude * \sin(period * \varphi + \pi)$$

By translating the first two rings along the z-axis, one can create a cylinder with rings of sinusoidal functions. Next, a vertical sine wave is generated based on the parameters of the rings. The starting and ending points of a vertical line are on the crossing points of the bottom and top circles and their baselines respectively. On this vertical wave, the degree $\varphi$ of the points varies with the z-coordinate as a sine function. Hence, the coordinate of every point on the vertical line is defined by

$$period = \frac{2\pi}{period\ length}$$

$$\Delta\varphi = amplitude * \sin(period * z)$$

$$\varphi = \Delta\varphi + \varphi_{baseline}$$

$$x = r * \cos(\varphi)$$

$$y = r * \sin(\varphi)$$

$$z = z_{baseline}$$

Where $\Delta\varphi$ is the diversion of the current point from the vertical baseline in terms of angle, and $period\ length$ is twice the distance between two rings. Then a second vertical sine wave with a phase shift of $2 * period\ length$ to the first wave is created. While the position of points on this line can be calculated using the same computations as above, the angle $\Delta\varphi$ contains a phase shift based on the first line:

$$\Delta\varphi = \ amplitude * \sin{(period * z \ + \ \pi)}$$

Furthermore, the angular distance $d$ between the baseline of the first and the second wave is

$$d = \frac{period\ length\ of\ sine\ wave\ circle}{2}$$

To obtain the final shape, all other vertical lines can be generated using the same method, while varying $\varphi_{baseline} \in [0, 2\pi]$ and adding phase shift to every second line.

***Auxetic cylinders with pinwheel meshes***

First, create a circle parallel to xy-plane. The x- and y-coordinates of all points on the circle vary with the angle $\varphi$, and the z-coordinates correspond to a sinusoidal function along the circumference. For a circle with radius $r$, the coordinates of each point on the circle are

$$x = r * \ \cos{(\varphi)}$$

$$y = r * \ \sin{(\varphi)}$$

$$\mathrm{z} = amplitude * \ \sin{(period * \ \varphi)}$$

$$period = \frac{2\pi}{period\ length}$$

where $\varphi \in [0,\ 2\pi],\ period\ length \in [0, 2\pi r]$. By translating the first ring along the z-axis, one can create a cylinder with parallel in-phase rings of sinusoidal functions.

Next, a vertical sine wave is generated based on the parameters of the rings. The starting and ending points of a vertical line are on the crossing points of the bottom and top circles and their baselines respectively. On this vertical wave, the degree $\varphi$ of the points varies with the z-coordinate as a sine function. Hence, the coordinate of every point on the vertical line is defined by

$$period = \frac{2\pi}{period\ length}$$

$$\Delta\varphi = \ amplitude * \sin{(period * z)}$$

$$\varphi = \ \Delta\varphi + \varphi_{baseline}$$

$$x = r * \ \cos{(\varphi)}$$

$$y = r * \sin(\varphi)$$

$$z = z_{baseline}$$

Where $\Delta\varphi$ is the diversion of the current point from the vertical baseline in terms of angle, and $period\ length$ is the distance between two rings. To obtain the final shape, all other vertical lines are generated using the same method, while varying $\varphi_{baseline} \in [0, 2\pi]$.

***Auxetic cylinders with vertically oriented arrowhead meshes***

First, create the vertical zigzag lines by defining the coordinates of all corner points. The corner points of every zigzag line can be described as points alternating on two sides of a base line. Hence, the x- and y coordinates of corner points are defined by

$$x = r * \cos(\varphi_{baseline} \pm amplitude)$$

$$y = r * \sin(\varphi_{baseline} \pm amplitude)$$

$Amplitude$ represents the distance between a corner point and the baseline in terms of angle, its sign alternates along the z-direction. Since all zigzag lines are parallel to each other, their corners lie on circles parallel to the xy-plane. The x- and y-coordinates of all points on a circle can be obtained by iterating around the circle with a certain angle step length. The total number of iterations necessary to create all points in a ring is:

$$n = \frac{2 * \pi}{angle\ step}$$

$Angle\ step \in [0, 2\pi]$ is the difference in angle between two neighboring baselines. To create the symmetry of the zigzag structure, $angle\ step$ is set as

$$angle\ step = 2 * amplitude$$

For $i \in [0,\ n-1]$, the x- and y-coordinates of all baselines are

$$x_{baseline} = r * \cos(i * angle\ step)$$

$$y_{baseline} = r * \sin(i * angle\ step)$$

Therefore, the x- and y-coordinates of all corner points on the bottom circle are

$$x = r * \cos(i * angle\ step + amplitude)$$

$$y = r * \sin(i * angle\ step + amplitude)$$

The second circle from the bottom has a certain phase shift to the first one, which can be introduced by the following coordinate formulas:

$$x = r * \cos(i * angle\ step - amplitude)$$

$$y = r * \sin(i * angle\ step - \ amplitude)$$

i.e. the sign of amplitude is opposite for two points with the same index in the first and second circle. Since the only constantly changing coordinate is the z-coordinate, all point circles can be created by translating the first two along the z-axis. After all the corner points of the vertical zigzag lines are defined, before generating the connecting beams, every beam between two corner points is first wrapped onto the cylinder coat using the wrapping method mentioned in re-entrant honeycomb 1.

Finally, to create the diagonal beams, every point on a vertical zigzag line is connected to two points on the neighboring zigzag lines.

***Auxetic cylinders with horizontally oriented arrowhead meshes***

First, create zigzag rings around the cylinder by determining coordinates of the corner points. The corner points of each zigzag ring are composed of two circles of points with a certain phase shift to each other. The total number of iterations necessary to create all points on one circle is:

$$n = \frac{2 * \pi}{angle\ step}$$

$Angle\ step$ is the difference in angle between two neighboring points in a circle. For $i \in [0,\ n-1]$, the x- and y-coordinates of the corner points on the first circle can be calculated by

$$x = r * \cos(i * angle\ step)$$

$$y = r * \cos(i * angle\ step)$$

Whereas the corner points on the second circle have the coordinates

$$x = r * \cos(i * angle\ step + \ phase\ shift)$$

$$y = r * \cos(i * angle\ step + \ phase\ shift)$$

To create a regular arrowhead structure, the phase shift is set to

$$phase\ shift = \frac{angle\ step}{2}$$

Then one can translate the coordinates from the first two circles vertically to create all circles along the z-direction. After defining all corner points of the zigzag rings, these can be connected to create the zigzag rings. Before generating the connecting beams, every beam between two corner points is wrapped onto the cylinder coat using the wrapping method mentioned in re-entrant honeycomb 1. Finally, diagonal beams were constructed between neighboring zigzag rings, i.e. every point in a ring is connected to two points on either its top or bottom neighbor in an alternating fashion.

***Auxetic mesh around the sphere***

First, create a cube with the edge length $a$ centered around the origin, and there are 4 edges parallel to x-axis, y-axis, or z-axis respectively. The corner coordinates of the cube are

$$(\pm a/2, \pm a/2, \pm a/2)$$

Then equidistant points are defined on every edge. Points on two opposite edges of a surface can be connected to construct the baselines for the periodic functions in later steps. All base lines between two parallel cube edges are parallel to each other and perpendicular to the cube edges. Then sine functions are created according to all baselines.

There are six kinds of sine functions used based on their plane propagation direction:

| Parallel plane / Propagation direction | yz-plane | xz-plane | xy-plane |
|---|---|---|---|
| x | | $(x_b, y_b, A * \sin(B * (\Delta x + C) + z_b)$ | $(x_b, A * \sin(B * (\Delta x + C) + y_b, z_b)$ |
| y | $(x_b, y_b, A * \sin(B * (\Delta y + C) + z_b)$ | | $(A * \sin(B * (\Delta y + C) + x_b, y_b, z_b)$ |
| z | $(x_b, A * \sin(B * (\Delta z + C) + y_b, z_b)$ | $(A * \sin(B * (\Delta z + C) + x_b, y_b, z_b)$ | |

$A$ is the amplitude of the sine functions. $B$ is the period coefficient

$$B = \frac{2 * \pi}{period\ length}$$

$$period\ length = \frac{a}{number\ of\ periods\ on\ a\ line}$$

$C$ is the phase shift which alters between 0 and $\pi$ for neighboring parallel lines. Additionally,

$$p_{baseline} = (x_b, y_b, z_b)$$

$$p_{start} = (x_s, y_s, z_s)$$

$$\Delta x = x_b - x_s$$

$$\Delta y = y_b - y_s$$

$$\Delta z = z_b - z_s$$

$p_{baseline}$ is a point on a baseline, $p_{start}$ is the starting point of that baseline. The coordinates of $p_{start}$ correspond to the predefined equidistant on cube edges. The resulting structure at this point is a cube with phase-shifted sinusoidal meshes on every surface.

Finally, all points are wrapped onto a sphere with radius $a/2$ using the following formula:

$$p_{sphere} = \frac{\vec{p}_{cube}}{|\vec{p}_{cube}|} * \frac{a}{2}$$

where $\vec{p}$ is the vector from the origin to point $p_{cube}$.

***Auxetic mesh wrapped around the heart***

First, a hemisphere covered by auxetic mesh is created following the same principle as the auxetic sphere above. After generating the cube with auxetic mesh, all points on the top half of the cube are removed. The gaps on the top edge of cube are connected by straight horizontal beams parallel to xy-plane. All points on the half cube can then be wrapped onto a hemisphere using the formula

$$p_{hemisphere} = \frac{\vec{p}_{cube}}{|\vec{p}_{cube}|} * \frac{a}{2}$$

Then a heart shape is placed into the hemisphere with a slight offset, i.e., the bottom of the heart intersects with the hemisphere, which ensures optimal fitting for the wrapping. Then every point on the auxetic hemisphere is wrapped onto the bottom of a heart using the built-in Hyperganic method "oNearestPointOnSurface", which computes the point on the heart surface with the smallest distance to a given point on the hemisphere lattice. Finally, this wrapping process is performed again between the new lattice and heart to optimize the result.

***Auxetic heart with wrapped auxetic patch***

First a curved auxetic sheet is created. Herein, one starts with a square 2D auxetic sheet on xy-plane. The edge length of the sheet is $l_{square}$. First, a circle with the diameter

$$d_{sheet} = l_{square}$$

is defined. In case the initial sheet is rectangular, the circle diameter is defined as

$$d_{sheet} = \min\{l_{square,x}, l_{square,y}\}$$

Where $l_{square,x}$ is the sheet dimension along x-axis and $l_{square,y}$ is the sheet dimension along y-axis. Then all points outside the defined circle are removed. Furthermore, the border line of the circle is create as part of the sheet. The result at this stage is a circular sheet.

Then all points need to be wrapped onto a curved surface with the given curvature $K$. This surface can be defined as a patch on a sphere with the radius

$$R_S = \sqrt{\frac{1}{K}}$$

The distance between the sphere center and the 2D sheet on the xy-plane is

$$H_C = \sqrt{{R_S}^2 - \left(\frac{d_{sheet}}{2}\right)^2}$$

A point $P$ on the 2D sheet is wrapped onto the sphere surface via translation along negative z-direction. The following relationship exists between the translation distance $|\Delta z|$, the distance $d(P, P_C)$ between $P$ and the sheet center $P_C$, $R_S$ and $H_C$:

$$(H_C + |\Delta z|)^2 + d(P, P_C)^2 = {R_S}^2$$

$$d(P, P_C) = \sqrt{\left(x_P - x_{P_c}\right)^2 + \left(y_P - y_{P_c}\right)^2 + \left(y_P - y_{P_c}\right)^2}$$

The translation distance $|\Delta z|$ can be obtained by reforming the equation above:

$$|\Delta z| = \sqrt{{R_S}^2 - d(P, P_C)^2} - H_C$$

Hence, the wrapped point $P_{projection}$ is

$$P_{projection} = P + (0, 0, -|\Delta z|)$$

All points on the 2D sheet on xy-plane can be wrapped using this principle. The resulting sheet is the partial surface of the sphere with negative z-coordinates.

The curvature of the sheet can be increased to fit the bottom of the heart by varying curvature $K$ in the calculations. After the sheet with the desired curvature is constructed, the heart is placed into the curved auxetic sheet such that they intersect slightly with each other, which improves the snapping result in the next step. Finally, every point on the auxetic sheet is wrapped onto the bottom of the heart using the built-in Hyperganic method "oNearestPointOnSurface", which computes the point on the heart surface with the smallest distance to a given point on the curved sheet.

***Perfusable channels***

All beams in this shape are aligned along the z-axis. The bottom part is constructed first, then the top part can be generated via mirroring. The curvature of the structure is created based on the hyperbolic sine function

$$z = \sinh(x)$$

within the interval $z \in [-10, 18]$. Then the curve is rotated 6 times around the center axis, i.e. for every point $p\ (x_p, y_p, z_p)$ with distance $d$ to the center axis on the curve, 6 additional points are created with coordinates

$$x = d * \cos(i * \frac{\pi}{4})$$

$$y = d * \sin(i * \frac{\pi}{4})$$

$$z = z_p$$

$$i \in [1, 6], i \in \mathbb{N}$$

The resulting curves form the bottom part of the channel structure with an six-fold symmetry around the center axis and the opening facing up.

Then the bottom structure is mirrored on a mirror plane parallel to the xy-plane with a pre-defined z-offset create the top structure. The two parts are then connected by vertical beams to form the completed shape. These channels are then removed from a cuboidal construct. In addition, a sphere with a pre-set offset is also removed from the cuboid to result in the perfusable construct with a hollow sphere in-between.

***Construct an alveolus***

First an STL file of an ellipsoid is inserted. Next, a foam structure is obtained inside the ellipsoid by applying the built-in Hyperganic method “SphericalFoamMap” to the sphere. This method creates small spheres with a certain distance to each other, these spheres are confined in the space of the ellipsoid. The Hyperganic method “Offset” is then used to enlarge all spheres (by factor 2), such that the spheres intersect with each other to build an entity. The Hyperganic method “Smoothen” (with factor 0.5) is used to smoothen the sharp intersection between spheres

***Geometry of a regular icosahedron***

First, the coordinates all 12 vertices need to be defined. The icosahedron in this case is centered around origin. First, only the 6 vertices lying on the top half of the icosahedron are determined, which consist of one vertex at the top and five vertices lying on a pentagon orthogonal to the z axis.

Start with a defined length $h$, which is the distance from every vertex of the icosahedron to its center. h is also the radius of the circumscribed sphere for the icosahedron, and the coordinate of the top point is $(0, 0, h)$. The relation between $h$ and the edge length $a$ is:

$$r = a * \sin(\frac{2\pi}{5})$$

Take $r = h$, then the edge length a of the icosahedron of the sphere is:

$$a = \frac{h}{\sin\left(\frac{2\pi}{5}\right)}$$

To determine the z-coordinate of the pentagon orthogonal to z-axis, one needs to compute the coordinate of one corner vertex on the pentagon. The geometric component needed for this computation is a triangular plane defined by the center of icosahedron, the top vertex, and one unknown corner vertex on the pentagon. Two sides of this triangle have the length h, and the third side is an edge on the icosahedron with length a. The following relation exists between h, a, and the angle $\varphi$ :

$$\sin\left(\frac{\varphi}{2}\right) = \frac{a/2}{h}$$

Since $h$ and $a$ are already determined from previous computations, solving the equation above gives the angle $\varphi$, which is approximately $63.43°$ for a regular icosahedron. The angle $\theta$ can be computed by $\theta = 90° - \varphi$. Then the orthogonal triangle defined by the center of icosahedron, the vertex on the pentagon and its wrapping onto the xy-plane is taken, and the following relations can be used to compute the vertical edge $l$ and horizontal edge $d$ of the triangle:

$$\sin(\theta) = \frac{l}{a}$$

$$\cos(\theta) = \frac{d}{a}$$

All vertices on the pentagon lie on a circle parallel to xy-plane with radius d. Therefore, for $i \in \{0, 1, 2, 3, 4\}$, each vertex has the coordinates

$$x_i = d * \cos\left(i * \frac{2\pi}{5}\right)$$

$$y_i = d * \sin\left(i * \frac{2\pi}{5}\right)$$

$$z_i = l$$

To construct the lower half of the icosahedron, which consists of a lower pentagonal plane and the bottom vertex of the whole icosahedron, one can use the 6 vertices obtained above. The bottom vertex is the mirrored point of its corresponding top vertex, and has the coordinates (0, 0, -h). The five vertices on the lower pentagon can be obtained by first mirroring the vertices on the upper pentagon by xy-plane to reverse the z-coordinates of all, then another mirroring procedure using the yz-plane. The resulting lower pentagon is rotated by $36°$ relative to the top pentagon.

Eventually, groups of 3 vertices are connected to create the 20 equilateral triangles that construct the icosahedron.

***Construct pentagons on icosahedron***
For every equilateral triangle on the icosahedron, its geometric center is connected with the midpoint of each edge.

***Construct hexagons on icosahedron***
First, every equilateral triangle is divided into 4 identical equilateral triangles by connecting the midpoints of all edges. This process can be repeated to further divide the triangles and create denser structures with smaller hexagons. Then, for every equilateral triangle, its geometric center is connected with the midpoint of each edge.

***Alveolus shape for printing***
This construction procedure is similar for both pentagonal- and hexagonal-shaped vessels. First, the color of the vessels is determined based on the coordinate of each point, i.e. all points on the left side are drawn in blue, and all points on the right in red. In the case of pentagonal vessels, to improve the quality of prints and avoid certain intersection with the top cylinder later, the pentagon on the top is enlarged by factor 2 to create a bigger opening.

Next, a thickness gradient is introduced to the vessels, such that the thickness decreases gradually from the left and right ends of the shape towards the middle. Given the maximum vessel thickness $t_{max}$ and the minimum vessel thickness $t_{min}$, the formula for the computation of thickness $t$ at point $P$ on the shape is

$$t_P = t_{min} + \left| (t_{max} - t_{min}) * \frac{|\Delta d_P|}{r/2} \right|$$

where $\Delta d_P$ is the horizontal distance between point $P$ and the middle of the shape, $r$ is the radius of the vessel shape.

Then every edge of the icosahedron is interpolated to create 11 equidistant intermediate points, i.e. 10 subbeams per edge, for better fitting in the following procedures. The entire shape is centered around the origin of the coordinate system. Then each point is wrapped onto the circumscribed sphere of the icosahedron by multiplying the sphere radius $r$ with the unit vector from the icosahedron center to the point.

$$P_{wrapping} = r * \frac{\vec{P}_{initial}}{|\vec{P}_{initial}|}$$

The result is a spherical vessel shape used for wrapping.

Then the alveolus is placed such that it’s concentric with the vessel shape. Then every beam on the vessel shape is interpolated into 10 subbeams and wrapped onto the alveolus using the Hyperganic method “oNearestPointOnSurface”, which determines the closest point on the alveolus to a givenn point on the vessels. This snapping procedure is then repeated once for to correct possible mismatches of the previous snapping step. The obtained shape constructs the final vessels.

To build the inlet, two points $P_+$ and $P_-$ with a deviation of $\sim \pm a/4$ from the middle line are chosen on one side of the shape. Then their branching point $P_{branch}$ is computed using the following formulas:

$$P_{middle} = P_+ + \frac{P_- - P_+}{2}$$

$$P_{branch} = \frac{l_{inlet}}{2} * \frac{\vec{P}_{middle}}{|\vec{P}_{middle}|}$$

$P_{middle}$ is the middle point between $P_+$ and $P_-$ and $l_{inlet}$ is the defined total length of the inlet. The outlet on the opposite side of the inlet is constructed following the exact same principle.

Then a cylinder is created and placed on top of the alveolus. Additionally, one can create a gap between the alveolus and the vessels by using a larger alveolus shape with the radius

$$r_{large} = r_{small} + gap\ distance$$

for wrapping of the vessels, then placing a smaller alveolus shape with radius $r_{small}$ concentrically for the final shape.

For volumetric printing, the finished alveolus shape with all components is subtracted from a cuboid, such that the alveolus is the empty part of the cuboid.

The dimensions of the vessels as well as the alveolus are tunable in the algorithm for the shapes. The ratio

$$alveolus\ diameter : vessel\ diameter : gap\ distance$$

for the printed object is $45 : 12 : 8$.

***Foamed shape***

First, the STL file of a solid shape was imported. Then the built-in Hyperganic function "VoronoiFoamMap" was applied on the shape, with defined intervals of cell dimension as well as wall thickness.

***Computational modeling of the structural deformation***

To evaluate the homogenized performance of the auxetic meshes a linear elastic material with Young's Modulus of $E = 10\ KPa$ and Poisson's ratio of $\nu = 0.4$ was assumed. The planar auxetic meshes were constrained with the roller support at the bottom of the mesh ($u_y = u_z = 0$) while applying a displacement boundary condition at the top of the mesh ($u_y = 10\ mm$) to induce 20% strain. The homogenized Poisson's ratio was computing by averaging a directional displacement component in an opposite direction:

$$\nu^* = -\frac{< u_x >_{x_{max}}}{L_x \cdot < \varepsilon_{yy} >}$$

Where $L_x$ is the length of the structure in the x direction. Cylindrical meshes were computed with the same material parameters as stated above. The boundary conditions were modified to reflect the cylindrical structure, in particular, an internal radial displacement was applied as:

$$u_x = u_0 \cdot \frac{x}{r}$$
$$u_y = u_0 \cdot \frac{y}{r}$$

To constrain the rigid body modes, the structure was fixed at the bottom. The homogenized Poisson's ratio was determined as:

$$\nu^* = -\frac{< u_z >_{z_{max}}}{L_z \cdot < \varepsilon_r >}$$

Where $< \varepsilon_r >$ refers to the applied radial strain.
